# Achieving pan-microbiome biological insights via the dbBact knowledge base

**DOI:** 10.1101/2022.02.27.482174

**Authors:** Amnon Amir, Eitan Ozel, Yael Haberman, Noam Shental

## Abstract

16S rRNA amplicon sequencing provides a relatively inexpensive culture-independent method for studying the microbial world. Although thousands of such studies have examined diverse habitats, it is difficult for researchers to use this vast trove of experiments when analyzing their findings and interpret them in a broader context. To bridge this gap, we introduce dbBact, an open wiki-like bacterial knowledge base. dbBact combines information from hundreds of studies across diverse habitats, creating a collaborative central repository where 16S rRNA amplicon sequence variants (ASVs) are manually extracted from each study and assigned multiple ontology-based terms. Using the >900 studies of dbBact, covering more than 1,400,000 associations between 345,000 ASVs and 6,500 ontology terms, we show how the dbBact statistical and programmatic pipeline can augment standard microbiome analysis. We use multiple examples to demonstrate how dbBact leads to formulating novel hypotheses regarding inter-host similarities, intra-host sources of bacteria, and commonalities across different diseases, and helps detect environmental sources and identify contaminants.

## Introduction

Bacteria play an important role in the Earth’s ecosystem, having a total biomass higher than that of all vertebrates and fish, second only to plants (1). The introduction of 16S rRNA amplicon sequencing as a means for molecular identification has enabled a culture-independent view of such ecosystems (2). Combined with massively parallel sequencing technologies and DNA barcodes (3), 16S rRNA sequencing provides relatively cheap and accurate microbial profiling. This, in turn, has led to a huge surge in the number of 16S rRNA studies examining microbial populations in habitats ranging across oceans (4), soil (5), plants (6), animals (7), and large cross-sectional human studies (8–10).

A severe limitation when combining insights from multiple microbiome studies is the complexity of the underlying bacterial populations, ranging from tens of different bacteria in a single saliva sample (11), to thousands in a single soil sample (12). In addition, although the total number of different bacteria is large (e.g., ∼300000 unique 16S rRNA sequences of length 90bp appear in the Earth Microbiome Project (12)), the number of taxonomic names for describing these bacteria is much smaller (about 3500-4000 unique genera and 20000 unique species names appear in NCBITAX (13) and in the Encyclopedia of Life (14)). Moreover, grouping bacteria in higher taxonomic levels may not always maintain the basic habitat properties of many bacteria (12). Therefore, reaching cross-study biological insights should preferably be based on 16S rRNA amplicon sequences rather than on taxonomy.

Recently, amplicon sequence variants (ASVs) derived using denoising methods such as Deblur (15), DADA2 (16) and UNOISE2 (17) have been introduced as an alternative to OTU picking for identifying bacteria in a given sample. Such denoising methods provide an objective identification of each bacterial sequence in the sample (i.e., independent of external databases or additional bacteria/samples in the experiment), as well as high sequence resolution (a single nucleotide difference in the sequenced region is identified as a different ASV). Therefore, ASVs may serve as cross-study identifiers for bacteria, i.e., a bacterium in different studies will result in the same ASV, even when the studies are processed separately and denoised using different methods (15,18).

In this paper, we introduce dbBact, a knowledge base for reaching cross-experiment biological insights. dbBact is based on manually collecting genotype-phenotype associations between ASVs and relevant conditions. For clarity, “reserved” dbBact words appear in italics: surveyed studies are referred to as *experiments*, stored ASVs are *sequences*, phenotypes are ontology-based *terms*, and genotype-phenotype associations are called *annotations*.

Currently, dbBact contains more than 900 *experiments* in various habitats, covering more than 1,400,000 associations between 345,000 *sequences* and 6500 ontology *terms*. For retrieval, dbBact provides two query types: a single *sequence*/FASTA file query, asking what is known about these *sequences* (Figure 1), and a query contrasting two FASTA files, searching for dbBact *terms* significantly enriched in either of the groups, analogously to gene ontology enrichment analysis (19,20) (Figure 2). By examining the ontology *terms* associated with each *sequence*, users can gain insights regarding the biology associated with ASVs of interest.

**Figure 1:**
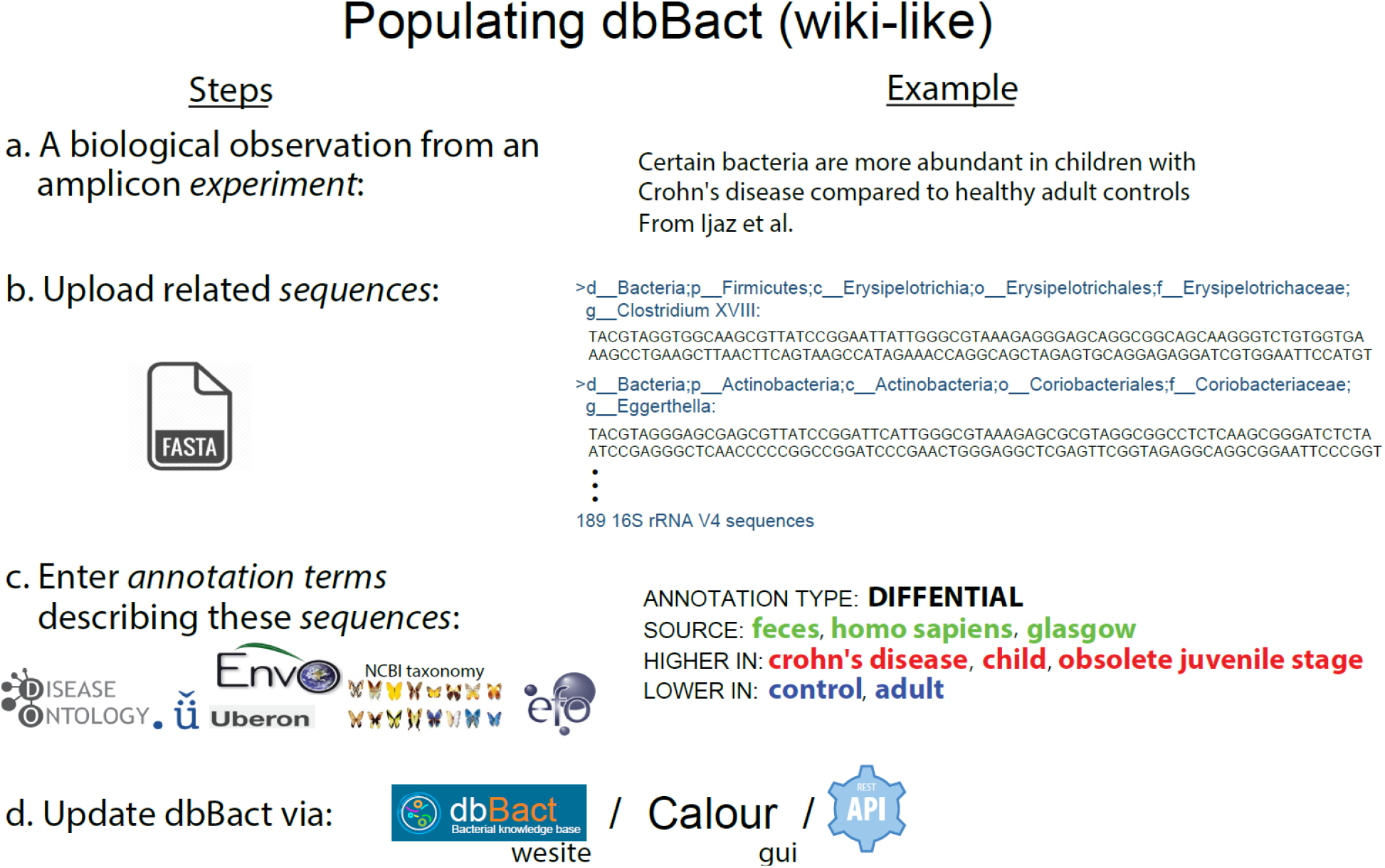
Adding entries to dbBact. Users add new entries in a wiki-like way, by uploading study results. **a**. For example, analyzing data from Ijaz et al. (37), we identified 189 ASVs that are more abundant in fecal samples of Scottish children with Crohn’s disease compared to healthy controls (see Methods section). **b**. These ASVs are uploaded as a FASTA file. **c**. Associations between ASVs and phenotypes are called *annotations*, which are created by assigning a set of ontology *terms* and predicates that characterize the context. The 189 *sequences* were *annotated* as “DIFFERENTIAL,” i.e., more abundant in children with Crohn’s disease (“HIGHER IN” *terms*), compared to healthy controls (“LOWER IN” *terms*). The general background *terms* common to both groups, i.e., “homo sapiens,” “feces” and “glasgow” are designated by “SOURCE.” *Terms* may be selected from several ontologies (e.g., DOID (33), ENVO (38,39), GAZ (40), UBERON (41), EFO (42), and NCBI Taxonomy (13)), allowing easy and precise *annotations*. **d**. Uploading *annotations* may be performed either through the dbBact website, dedicated clients (i.e., Calour (36)) or by REST-API. For clarity, the following nomenclature holds throughout the manuscript where “reserved” words appear in italics (e.g., *experiment, sequence, annotation, term*), predicates appear in all caps (e.g., “HIGHER IN,” “LOWER IN,” “SOURCE”), and specific *term* names follow the ontology convention of being lower case (e.g., “homo sapiens”).

**Figure 2:**
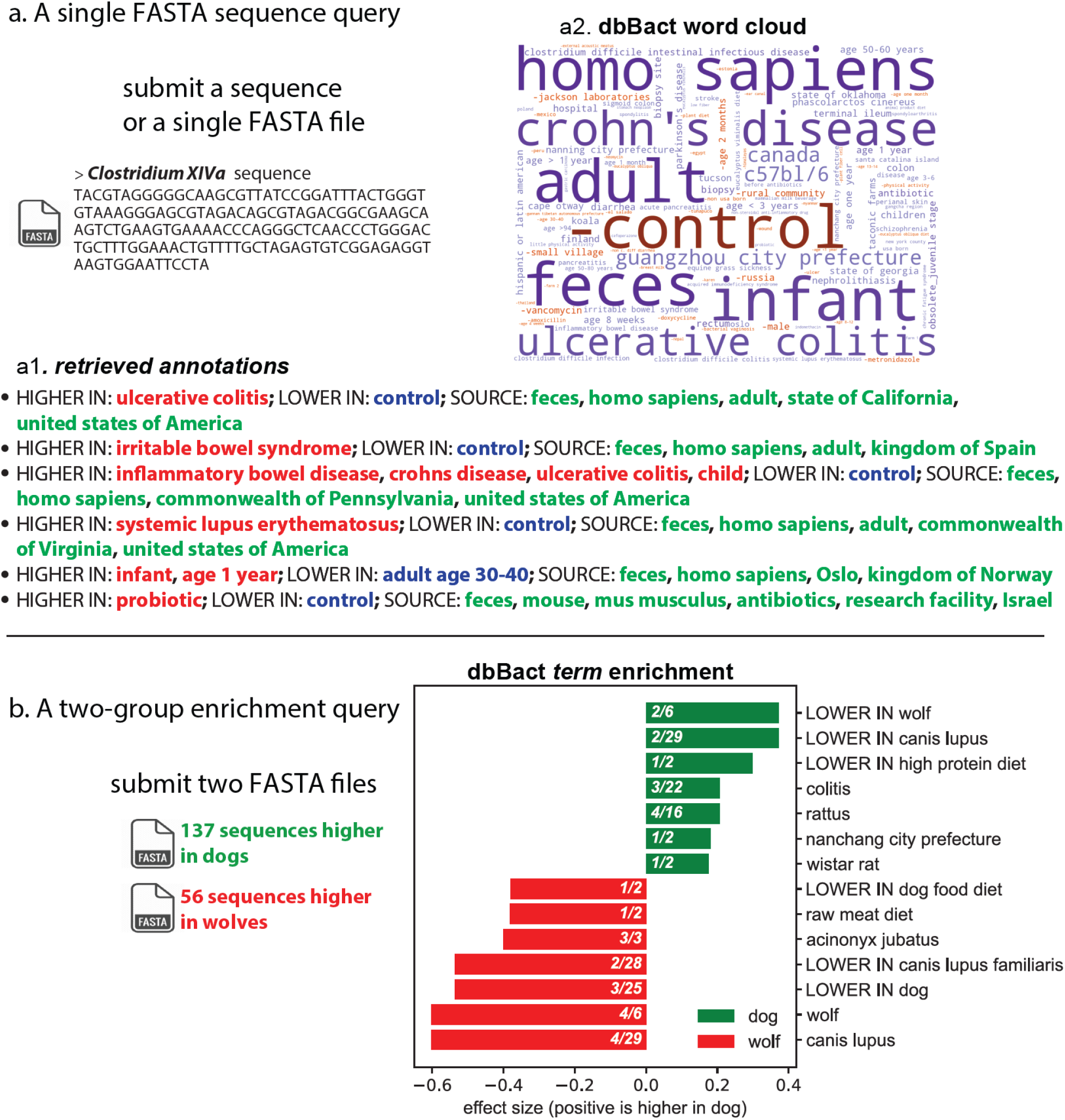
Two basic query types: **a**. Uploading a FASTA file of *sequences* results in a list of the most relevant *annotations* containing these *sequences*, and a “word cloud” of best matching *terms*. In this example, a V4 *sequence* of *Clostridium XIVa*, which is highly abundant in fecal samples of chronic fatigue syndrome patients (CSF) (Giloteaux et al., 2016), was submitted. Panel **a1** provides representative *annotations* containing the query sequence (the full list of ∼150 *annotations* appears in Supplementary File 3). dbBact found this *sequence* to be higher in the disease group than in healthy controls in several studies (ulcerative colitis, irritable bowel disease, and lupus), and in antibiotic-treated mice supplemented with probiotics (last *annotation* arising from (43)). Panel **a2** displays the word cloud summarizing the *terms* associated with the query *sequence*, where size corresponds to a *term*’s F1 score, while color designates the associated predicate (blue for “SOURCE”/”HIGHER IN” *terms*, and red color preceded by a minus sign corresponds to “LOWER IN” *terms*). Hence, this *Clostridium XIVa* query *sequence* is associated with human feces in dysbiosis states of “crohn’s disease,” “ulcerative colitis,” “diarrhea,” and “c. difficile infection” (a full list of F1 scores per term appears in Supplementary File S7). **b**. By contrasting two groups of *sequences*, dbBact identifies enriched *terms* characterizing each group. For example, 137 and 56 *sequences* were submitted, corresponding to differentially abundant *sequences* higher in fecal samples from domestic dogs and wolves living in zoos, respectively (data from (44)). Bar lengths show the normalized rank-mean difference for the top significantly enriched *terms* in the dog and wolf *sequences* (green and red bars, respectively). *Term* enrichment is based on a non-parametric rank mean test with FDR<0.1 using dsFDR (see the term enrichment analysis section in Methods). The numbers in the bar of each *term* correspond to the number of dbBact *experiments* in which the *term* differs significantly between the two *sequence* groups (numerator) and the total of dbBact *experiments* containing the *term* (denominator). *Sequences* that were more abundant in the wolf group are enriched in *terms* related to wolf, meat diet, and cheetah (*Acinonyx jubatus*).

dbBact differs from other microbial databases in several aspects: (a) *Manual annotation*: dbBact phenotype-genotype associations are extracted using manual analysis, in contrast to microbial data repositories, such as SRA/EBI, Qiita (21), HMP DACC (10,22), MGnify (23), FoodmicrobioNet (24), and redBIOM (25), which provide raw experimental data and metadata. In dbBact, the human expert understands the experimental setting and identifies abundant bacteria in different study groups, detects contaminants, etc., and these associations are uploaded. (b) *Unlimited scope*: dbBact accepts studies across all habitats, unlike databases that are highly limited in scope, e.g., human-disease-focused databases (MicroPhenoDB (26), gutMDisorder (27), Disbiome database (28), Peryton (29), and BugSigDB (30)), or other context-specific databases (database of the healthy mouse microbiome (31) or the sponge microbiome project (32)). (c) *Volume and potential growth*: to date, the number of studies in dbBact is ∼40% higher than in Qiita, and the number of ASVs is comparable to those in the Earth Microbiome Project. New studies are continually added by the dbBact team. Additionally, as a wiki-like database, we encourage the microbiome community to contribute to dbBact. (d) *Structured genotype/phenotype search*: observations are uploaded at the ASV level, allowing queries of specific *sequence*s and sub*sequence*s. In addition, as phenotypes are designated by *terms* derived from multiple ontologies, subsequent querying allows for “cross-sectioning” of the data. For example, *sequences* associated with Crohn’s disease and ulcerative colitis both originate from the DOID ontology (33), and will be recalled when querying their “parent” *term*, “inflammatory bowel disease.” (e) *Harmonizing studies performed using different variable regions*: as uploaded studies may be *sequence*d using different 16S rRNA variable regions, stored *sequence*s are “linked” through the SILVA database of full-length 16S rRNA genes (34), facilitating cross-region queries. For example, when submitting a query *sequence* from V1-V2, dbBact seeks the matching full-length 16S rRNA genes in SILVA, then extracts their V4 region, and subsequently retrieves relevant *annotation*s. (f) *Data analysis*: dbBact provides a set of statistical tools for analyzing new studies and for generating novel biological hypotheses.

In the following sections, we present the current scope and comprehensiveness of dbBact, as well as demonstrate, using multiple examples, how dbBact may be incorporated into standard microbiome analysis pipelines, thus providing novel hypotheses.

dbBact can be accessed using its website (http://dbbact.org), plugins for Qiime2 (35) and Calour (36), and programmatically using the dbBact REST-API interface.

## Results

### dbBact: Scope and comprehensiveness

dbBact release 2022.07.01 contains approximately 345,000 unique bacterial amplicon *sequences*, an amount that is on par with the 300,000 sequences observed by the Earth Microbiome Project (12). *Sequences* arise from over 900 unique *experiments*, i.e., studies from which observations were added (Figure 3a). Over 7000 dbBact *annotations* associate these *sequences* with various phenotypes using ontology derived *terms*. As each annotation typically includes many *sequences*, this results in over 1,400,000 unique genotype-phenotype associations.

**Figure 3:**
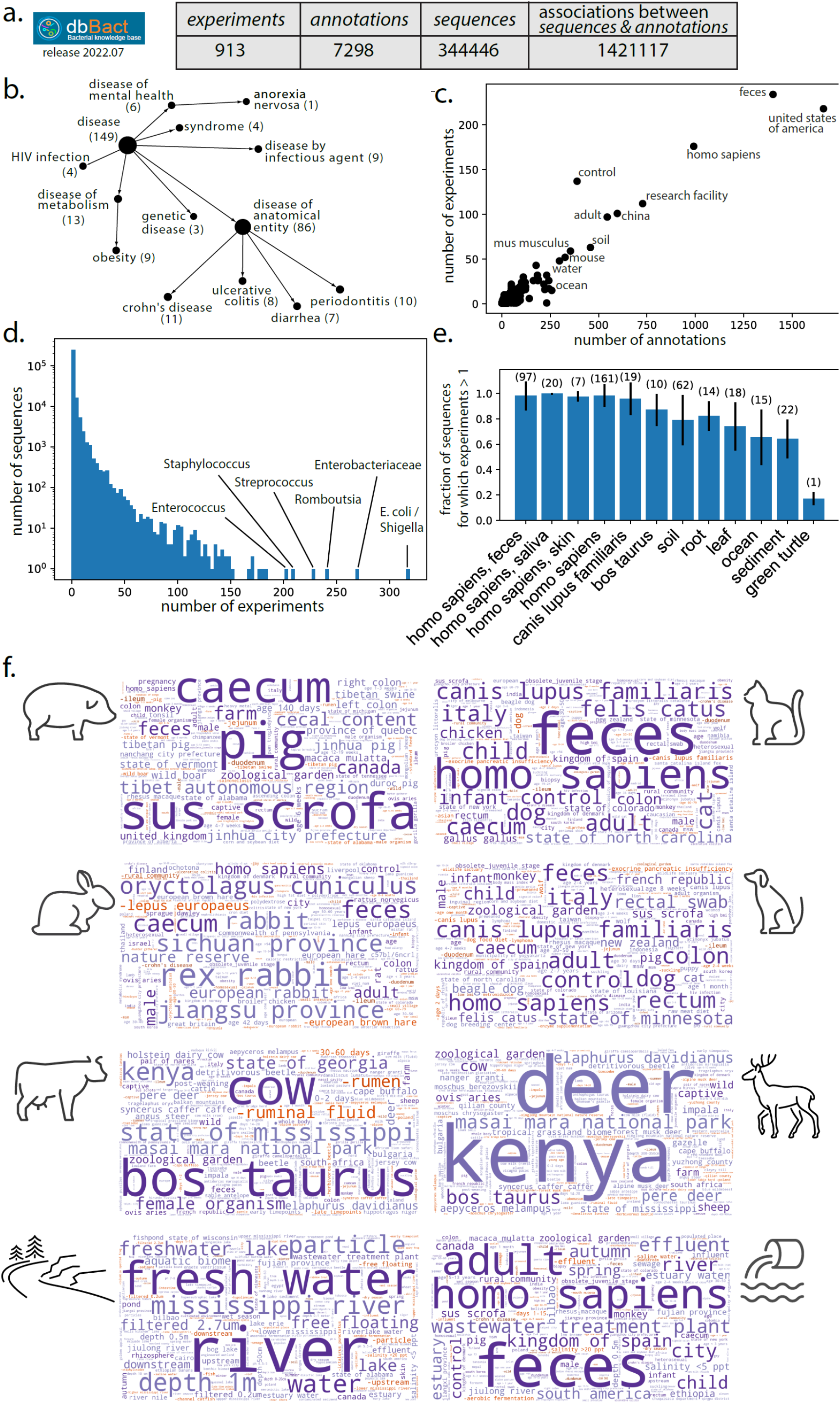
Knowledge base scope and comprehensiveness. **a**. Scope of dbBact release 2021.05 (used for the analysis presented in this paper). **b**. The number of *experiments* for representative disease categories based on the DOID ontology. **c**. Scatter plot of the total number of *annotations* and *experiments* in which each dbBact *term* appears. **d**. Histogram of the number of *experiments* in which each dbBact *sequence* appears. **e**. Knowledge base comprehensiveness. The fraction of “COMMON” *sequences* from each *experiment* that have been annotated in additional *experiments* is shown for various *terms*. The number of experiments containing the *term* is designated above each bar. **f**. Comprehensiveness in a source tracking task. *Sequences* from eight sample types from Hägglund et al. were blindly submitted to dbBact. Their word clouds clearly display the sources of the samples (shown by the matched cartoon). *Term* sizes correspond to the F1 score of each *term*, combined for all *sequences* present in > 0.3 of the samples (for each sample type).

### General statistics of dbBact

The >900 dbBact *experiments* cover a wide range of habitats (Figure S1a), geographic regions (Figure S1b), plant and animal hosts (Figure S1c), human body sites (Figure S1d), and human diseases (Figure 3b). For example, 149 *experiments* cover diseases, of which 86 are of an anatomical entity (e.g., Crohn’s disease or ulcerative colitis), and seven more are defined as metabolic diseases. The most abundant dbBact *terms* are “united states of america,” “homo sapiens,” and “feces,” each appearing in over 1000 *annotations* arising from more than 150 different *experiments*. Most of the other *terms* appear in less than twenty *experiments* (Figure 3c). The most prevalent bacterial *sequence* is *E. coli*, appearing in over 900 *annotations* from over 300 *experiments* (Figure 3d). Although this could reflect the universality of *E. coli* in various habitats, it may also be due to potential contaminations (45), a reason that may also explain the high prevalence of *Staphylococcus* (appearing in over 200 dbBact *experiments*). The number of *experiments* per *sequence* follows a power law distribution, with a majority of *sequences* appearing in a single *experiment*, yet over 80,000 *sequences* were observed in more than one *experiment* and 7000 *sequences* appeared in at least ten *experiments* (Figure 3d).

dbBact allows the upload of *sequences* from several commonly used regions (V1-V2, V3-V4 or V4; see Table S2 for a list of primers). Upon upload, *sequences* from different regions are “linked” through their full-length 16S rRNA sequence in the SILVA database (34) (see Inter-region querying section in Methods). When submitting query *sequences* from one region, dbBact retrieves all *annotations* containing the corresponding *sequences* across all regions (including, naturally, the region from which the query was provided). To demonstrate the usefulness of such “linking,” Figure S2 provides several examples of V1-V2 and V3-V4 *sequence* queries that are successfully characterized based solely on “linked” V4 *sequences*.

### Comprehensiveness of dbBact

#### Intra-dbBact estimates

To estimate the comprehensiveness of dbBact, we tested how many bacterial *sequences* typical of a specific environment (e.g., human feces) have *annotations* arising from more than one *experiment*. We selected *sequences* having an *annotation* of type “COMMON” (i.e., present in more than half of the samples in an *experiment*) for each of several *terms*, and measured the fraction of these *sequences* that have *annotations* from another dbBact *experiment* (Figure 3e). For example, there are 97 *experiments* having a “COMMON IN homo sapiens feces” *annotation*. Iterating over each of these *annotations*, about 98% of the associated *sequences* appear in more than one *experiment*. Hence, fecal bacteria are already well covered by dbBact. A similar level of “coverage” occurs for several other human-related *terms* and for dogs, where almost all *sequences* were observed in more than a single *experiment*. Regarding the *terms* “cow,” “soil,” “root,” and “leaf,” about 80% of the *sequences* appear in more than a single *experiment*, whereas the “coverage” of “green turtle” is much lower, indicating that additional experiments are required to capture its full bacterial diversity.

#### Out-of-sample comprehensiveness

As another example of comprehensiveness, dbBact was tested in a source tracking task, i.e., identifying the host or niche of a sample based on its bacterial composition. Hägglund et al. collected samples from either sewage influent or from freshwater, as well as feces sampled from several animals (rabbit, cat, wild boar, dog, cow and deer), aiming to find unique bacterial footprints of each source (46). We used all *sequences* present in more than 1/3 of the samples from each group as queries to dbBact resulting in word clouds describing each sample group (Figure 3f). In almost all cases, the notable *terms* in each word cloud were indicative of the sources of the samples, e.g., *sus scrofa* for the wild boar fecal sample, allowing accurate source tracking. The only exception was cat fecal samples, which were detected as a combination of cat, dog, and human, probably because of the small number of cat fecal samples present in the current dbBact release.

### The advantage of sequence-based associations

Results of 16S rRNA profiling experiments comprise a list of ASVs found in each sample and their abundances. Corroboration of these results with other microbiome studies is typically performed by searching published studies mentioning the taxonomy of these sequences. In many cases, however, such text-based mining may be limited because of constraints in taxonomic assignment. First, taxonomy is far from being full, e.g., species-level assignment is missing for about 80% of 16S rRNA sequences in Greengenes (47), and about 35% of the Greengenes sequences lack a genus assignment (48). Second, in many cases the same assigned taxonomy may be associated with vastly different phenotypes. As observed by the Earth Microbiome Project, bacteria of the same genus may be present in vastly different habitats, whereas specific sequences are associated with a certain habitat (12). This phenomenon underscores the importance of sequence-based association as provided by dbBact. For example, both *sequences* in Figure 4a belong to the genus *Blautia*, hence taxonomy-based associations may conclude that they play similar “roles” and are associated with the same phenotype. But querying dbBact with each of these two *sequences* results in a strikingly different picture, which we refer to as a “good” and “bad” *Blautia*. The “good” *Blautia* is more abundant in healthy controls than in patients of type 1 diabetes (T1D), Crohns’ disease (CD), inflammatory bowel disease (IBD), diarrhea, and kidney stones (Figure 4a), whereas the “bad” *Blautia* is more prevalent in patients suffering from IBD, CD and ulcerative colitis (Figure 4b).

**Figure 4:**
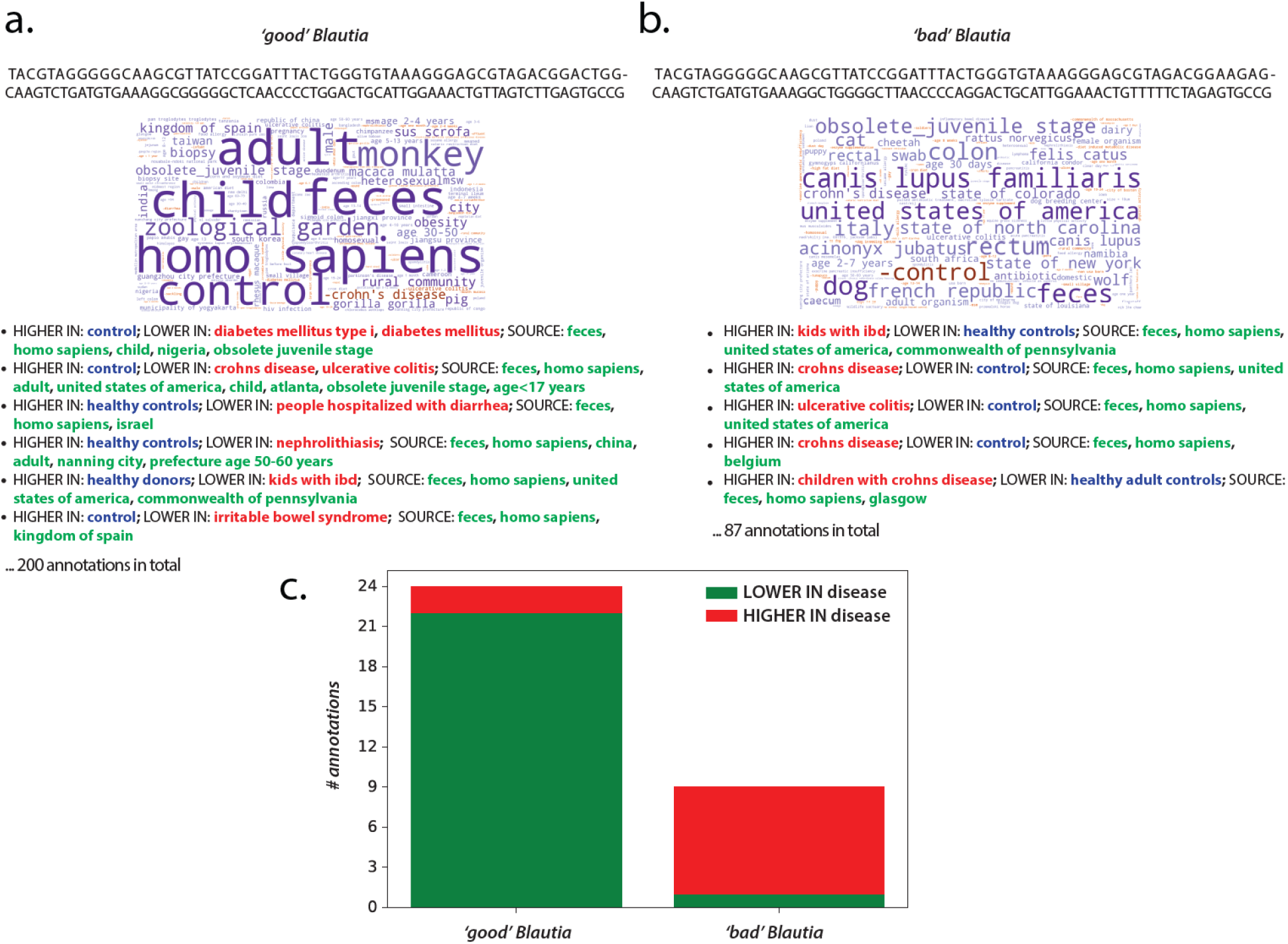
Taxonomy may be misleading. **a**. Two *sequences* of the genus *Blautia*, that differ by nine bases over the 150bp Illumina read of the 16S rRNA V4 region are associated with opposite phenotypes, as discovered by dbBact. The two word clouds and annotations for each *sequence*, display “opposite” associations with disease. The left *sequence* is more prevalent in healthy subjects (“good” *Blautia*), whereas the other is highly abundant in a series of disease-related *annotations*. Such differences can be traced through dbBact, but are completely missed by a taxonomy-based analysis. **b**. The number of disease-related *annotations* for the two *Blautia sequences* across dbBact displays an opposite trend of being low and high in disease, for the “good” and “bad” *Blautia*, respectively. The total number of *annotations* in dbBact 2021.05 associated with the “good” and “bad” *Blautia sequences* is 377 and 124, respectively.

Collecting all “disease” related dbBact annotations shows that the “bad” *Blautia* is “HIGHER IN” in the disease group (compared to controls) in 8/9 disease *annotations* associated with it, whereas the “good” *Blautia* is “LOWER IN” in the disease group (compared to controls) in 22/24 disease *annotations* (Figure 4c). Therefore, sequence-based analysis provides a solid genotype-to-phenotype association compared to taxonomy-based associations.

### dbBact provides a pan-microbiome view: Detailed example

dbBact may add another layer to data analysis in microbiome studies by identifying commonalities between different conditions and diseases, generating novel biological hypotheses. To demonstrate such a pan-microbiome analysis, we use data from a study comparing subjects consuming an American diet to a calorie restricted diet (49), and demonstrate the use of dbBact *term* enrichment. Fecal samples from two groups of lean individuals (BMI<25) who followed either an American diet (AMER) or a caloric restriction diet (CR) were selected. Standard analysis with FDR set to 0.1 (see “standard analysis” section in Methods) identified 28 and 141 bacterial *sequences* significantly more abundant in AMER and CR cohorts, respectively (Figure 3a). For clarity, we refer to these groups of *sequences* as S-AMER and S-CR, respectively. Figure 3b shows the internal “transformation” performed by dbBact from a heatmap of bacterial abundances to a heatmap of association scores of *terms* for each *sequence* (columns in Figure 3a-b are aligned and correspond to the same *sequences*). For example, the *term* “high BMI,” appears in almost all S-CR *sequences*, while it is almost absent in S-AMER *sequences*. These association scores in Figure 3b are then used as input to a non-parametric differential abundance test (see Methods section “statistical analysis in dbBact”), identifying *terms* significantly enriched in each of the two sequence groups (Figure 3c). Results indicate that *sequences* in the S-CR group are associated with *terms* related to low BMI (“low bmi,” “LOWER IN high bmi”) and with rural/undeveloped habitats (“LOWER IN united states of america,” “small village”) (see Supplementary File 4 for the full list of enriched terms). By contrast, bacteria from the S-AMER group have a significantly higher number of *annotations* related to high BMI (“LOWER IN low bmi,” “high bmi”) and urban/modernized habitats (“state of oklahoma,” “LOWER IN rural community,” “LOWER IN small village”).

Thus, although participants from both diet groups were lean, certain aspects of the underlying microbiome were associated with high and low BMI bacteria, for AMER and CR, respectively. Additionally, bacteria enriched in CR vs. AMER tend to be associated with rural/undeveloped habitats, which may indicate an adaptation of some bacteria found in rural communities to a low-calorie/higher vegetable diet content.

To further confirm the relationship between diet and BMI, we collected all *sequences* across dbBact having a “high bmi” annotation, resulting in 319 *sequences*. The overlap between these *sequences* and the S-AMER and S-CR groups is shown in Figure 3d (left). Although 93% (26/28) of S-AMER *sequences* overlap with “high bmi”-associated *sequences*, the overlap of S-CR *sequences* is 11% (15/141), i.e., a much larger fraction of S-AMER *sequences* is associated with high BMI. An analogous Venn diagram for the *term* “low bmi” displays an overlap of 54% (15/28) and 78% (110/141) of S-AMER and S-CRON bacteria with “low bmi”-associated bacteria in dbBact, respectively (Figure 3d left). As participants from both CR and AMER groups were lean (BMI<25), one may hypothesize that the effect of BMI on the microbial composition, observed in various studies, is due to dietary differences rather than the high BMI phenotype.

#### Remark regarding spurious/irrelevant terms

dbBact release 2022.07.01 contains *annotations* of approximately 6500 unique *terms*, some appearing only in a few *experiments*. As a result, word clouds and bar plots may often include seemingly odd *terms*. For example, the *term* “state of oklahoma” in Figure 5d is significantly enriched in S-AMER, a fact that seems implausible. This *term* appears only in two dbBact *experiments*, one of which compared a rural community in Peru to an urban community in Oklahoma (50). Hence, *annotations* from this *experiment* mentioned the *term* “state of oklahoma” together with more relevant *terms* (e.g., “rural community”) which, in turn, caused its inclusion. As dbBact continues to grow, such “transient” irrelevant inductions are expected to diminish.

**Figure 5:**
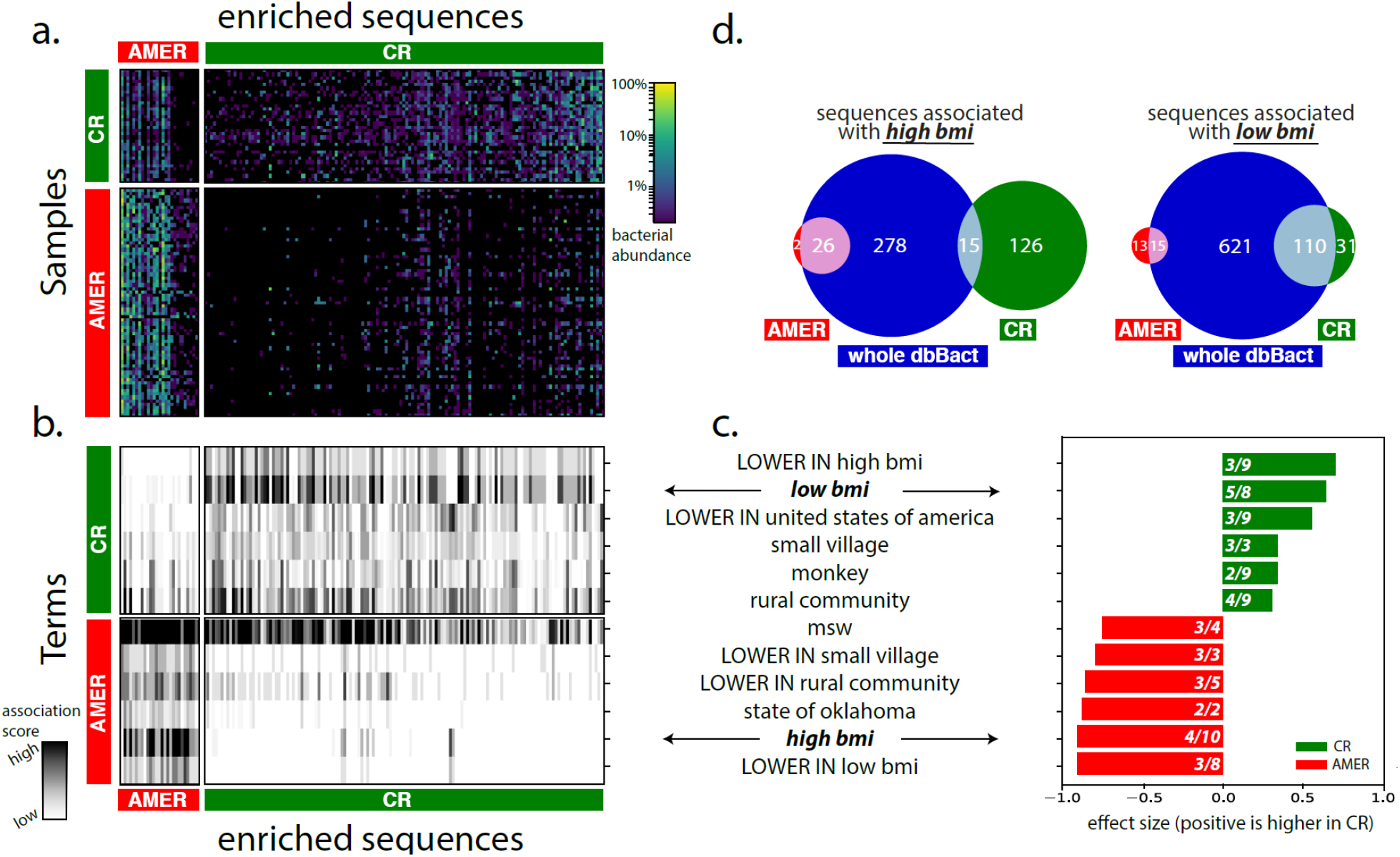
Linking caloric restriction associated bacteria to other phenotypes. **a**. Heatmap displaying bacterial abundances across fecal samples (rows) of low BMI individuals (BMI<25) practicing either a caloric restriction diet (CR, n=33) or an American diet (AMER, n=66), over a set of *sequences* (columns) that are significantly higher in either group. A differential abundance test (rank-mean test with dsFDR=0.1 multiple hypothesis correction) identified 136 bacteria higher in the CR group (S-CR) and 27 bacteria higher in the AMER group (S-AMER). **b**. dbBact *terms* (rows) enriched in the *sequences* appearing in panel **a** (columns in panels **a** and **b** are aligned). Heatmap values indicate the *term* score for each bacterium. *Terms* were identified using a non-parametric rank mean difference test with dsFDR=0.1 (top 6 terms for each direction are shown; see Supplementary File 4 for full list of enriched *terms*). **c**. Summary of the top enriched *terms* in the CR and AMER diets (green and red bars, respectively). Bar length and numbers are as in Figure 2. **d**. Venn diagrams of dbBact *annotations* related to the *terms* “low bmi” (right) and “high bmi” (left). Green and red circles indicate the number of *sequences* associated with the *term* in the CR and AMER diets, respectively; the blue circle indicates the number of such *sequences* across dbBact as a whole. The intersections of “low bmi” bacteria with the CR group are significantly higher (p=7E-5, using two-sided Fisher’s exact test), confirming the association. Similarly, the intersection of “high bmi” annotated *sequences* across dbBact with the AMER group is significantly higher than that with the CR group (p=3E-17, using two-sided Fisher’s exact test).

### Integrating dbBact into microbiome analysis pipelines allows generating novel biological hypotheses

To demonstrate how dbBact may be incorporated into microbiome analysis pipelines, we chose 16 dbBact *experiments*, excluded them from dbBact, then analyzed their results in the same way researchers would look at their own studies. In each of these studies, dbBact provided novel hypotheses that did not appear in the original paper and could not be formulated by standard methods.

The *experiments* presented here were chosen almost arbitrarily to provide examples from different habitats and niches: the human host, animals, and environmental samples (Figure 6a). dbBact-derived hypotheses may be divided into several “types,” as follows.

**Figure 6:**
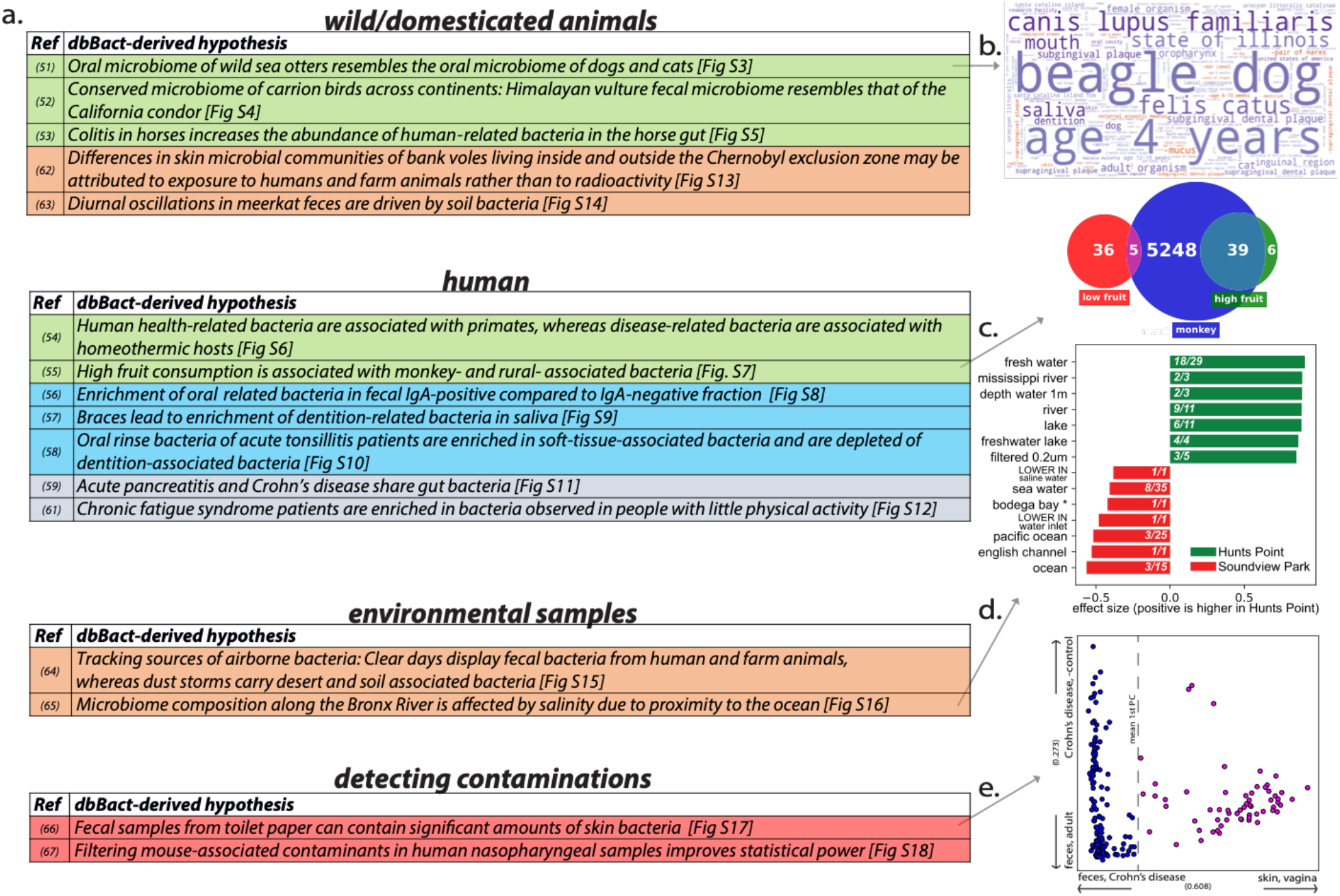
dbBact leads to novel biological hypotheses. **a**. Summary of biological hypotheses derived from dbBact-based analysis of published studies. Details of each analysis are given in the corresponding Supplementary Results section. Row colors correspond to hypothesis “type” (inter-host similarities – green; intra-host similarities – blue; inter-disease similarities – gray; environmental sources – brown; contamination detection – red). **b-e**. Analysis results related to conclusions shown in panel a. **b**. dbBact *term* word cloud for *sequences* found in sea otter oral samples shows resemblance to dogs’ and cats’ samples. **c**. Venn diagram showing number of dbBact *sequences* associated with the *term* “monkey” across dbBact (blue), and their intersection with *sequences* found in individuals from the American Gut study, who consume a high (green) and low (red) number of fruits per week. *Sequences* in the high-fruit consumption group are significantly more associated with the *term* “monkey” (Fisher’s exact test p-value < 0.00001). **d**. dbBact *term* enrichment comparing water samples collected in Hunts Point and Soundview Park, along the Bronx River in New York. *Sequences* higher in Hunts Point (green, located upstream) show significant fresh-water-related *term* enrichment (dsFDR=0.1). **e**. *Term*-based PCA of fecal samples of one individual collected daily for one year. The first principal component is the “feces-skin” axis, where higher values correspond to “skin” (see Methods for details). The values of a subset of samples, shown in magenta, is high, indicating possible skin-derived contamination in these fecal samples.

#### Detecting inter-host similarities

dbBact can identify unexpected similarities in microbial populations across hosts. (*i*) For example, when examining the oral microbiome of wild sea otters (51), dbBact indicates a high similarity to the microbiome of the oral cavity of dogs and cats (Figure 6b and Figure S3). (*ii*) In another example, fecal bacteria of Himalayan Griffons (52) are found to be similar to those of another carrion feeder, the California Condor (Figure S4). (*iii*) Such inter-host similarities are also observed for disease-related bacteria. Examining bacteria in colitis in horses (53), dbBact detects an enrichment of human-associated bacteria, indicating a possible colonization by bacteria that are less host-specific (Figure S5). (*iv*) Another recent meta-analysis of various human diseases identified shared disease-related bacteria in multiple diseases (54). When examining non-human-related *annotations*, dbBact finds these bacteria to be enriched in non-primate, homeothermic animals (mouse, horse, rat, chicken). By contrast, health-related bacteria found in this study are enriched in monkey-associated *terms* (Figure S6). This may indicate the disappearance of host-specific bacteria in multiple diseases, together with the appearance of more generalist bacteria. (*v*) A similar enrichment in monkey-associated bacteria and rural-community related *terms* is observed in individuals from the American gut project (55) who report high consumption of fruits, compared to those reporting low consumption (Figure 6c and Figure S7).

#### Detecting intra-host similarities

dbBact can identify similarities within hosts. (*i*) For example, Scheithauer et al. (56) profiled the bacteria detected in the IgA-positive and IgA-negative fractions of fecal samples. dbBact-based analysis shows that the IgA-positive fraction is enriched in oral related *terms*, indicating a possible contribution of oral IgA to bacterial antibody coating (Figure S8). (*ii*) In another study (57), dbBact finds an enrichment in dentition-related *terms* in an oral rinse of adolescents with braces compared to an enrichment in soft-tissue-associated bacteria in those that do not wear braces (Figure S9). (*iii*) Such soft-tissue-associated bacteria are also observed when analyzing Yeoh et al. data (58) of tonsilitis patients (Figure S10).

#### Detecting inter-disease similarity

(*i*) Zhu et al. compared the fecal microbiome of acute pancreatitis patients with that of healthy controls (59). dbBact-based analysis hints at a common gut response between pancreatitis and diarrhea, and Crohn’s disease, i.e., a phenomenon of general dysbiosis formerly suggested by Duvallet et al. (60) (Figure S11). (*ii*) Giloteaux et al. (61) compared fecal samples of chronic fatigue syndrome patients with those of healthy controls. dbBact finds shared *sequences* between these patients and individuals who do little physical activity (Figure S12).

#### Detecting environmental sources

dbBact can detect the sources of bacterial communities. (*i*) For example, Lavrinienko et al. collected skin swabs of bank voles inside the uninhabited Chernobyl exclusion zone and outside the contaminated region in the outskirts of Kyiv (62). dbBact-based analysis shows an overrepresentation of soil- and plant-related bacteria inside the exclusion zone, while skin bacteria of bank voles near Kyiv were enriched in human and farm animal *terms*. This leads to the hypothesis that the difference between the two sample groups is due to contact with humans and farm animals rather than to exposure to radioactivity (Figure S13). (*ii*) Similarly, Risely et al. (63) observed strong diurnal oscillations in the microbiome composition of South African wild meerkats’ fecal samples. dbBact-based analysis indicates that this effect is driven by a large number of soil/rhizosphere-related bacteria appearing in the afternoon fecal samples (Figure S14). (*iii*) dbBact analysis of air samples taken by Gat et al. (64) during clear days in Israel shows human farming as a source of air bacteria, compared to samples taken during a dust storm, which display desert and soil-associated bacteria (Figure S15). Hence, fecal bacteria from human and farm animals are airborne during ambient weather conditions, whereas dust storms bring over desert and soil associated bacteria. (*iv*) Finally, analysis of river water samples in two locations near the Bronx River estuary (65) shows that the difference in bacterial communities in these two locations is partially explained by ocean vs. freshwater bacteria, probably related to the salinity levels in the two sample locations (Figure 6d and Figure S16).

#### Contamination detection

dbBact allows the straightforward detection of potential contaminants. Each bacterium in a study may be assigned its best fitting dbBact *term*, thus “awkward” bacteria may be detected and discarded from downstream analysis. (*i*) Caporaso et al. (66) followed the oral, skin, and fecal microbiome of an individual using daily samples for a year. dbBact-based analysis detected a group of skin-associated *sequences* in a subset of fecal samples, indicating a potential contamination (Figure 6e and Figure S17). (*ii*) Similarly, in a dataset of infant nasopharyngeal samples (67), we observed a cluster of mouse-associated *sequences* (Figure S18), which may be attributed to a contamination or to low biomass kit-related bacteria. As these mouse-associated sequences are evenly spread across the sample types, they did not introduce a systemic bias in the authors’ results. But removal of the *sequences* before downstream analysis reduces inter-sample noise and increases the statistical power (Figure S18c,d).

## Discussion

dbBact integrates 16S rRNA microbiome studies into a collaborative, coherent body of knowledge that facilitates pan-microbiome analysis of new studies using a rigorous statistical and algorithmic framework.

An important advantage of dbBact, compared to standard meta-analysis methods, is that the latter may suffer from the “streetlight effect” (68). For example, when examining the effect of fruit consumption in the American Gut experiment (Fig 6a), one might consider including other diet-related studies in the meta-analysis. But this would miss the link between high fruit consumption and primate-associated bacteria. dbBact retrieves *annotations* from a wide range of sample types and habitats, providing additional and potentially unexpected insights into the biological roles of bacteria.

*Terms* in dbBact *annotations* are based on ontologies, providing a common language for phenotype description. The tree structure of ontologies facilitates the discovery of commonalities between bacteria in studies conducted under similar, albeit not identical, conditions. For example, data from Crohn’s disease and ulcerative colitis *experiments* may be combined based on their ontological “parent” *term* “inflammatory bowel disease.” Moreover, many “cross-sectional” questions may be asked and possibly answered using dbBact. For instance, what *terms* are similar with respect to their bacteria (e.g., are dogs more similar to cats or to wolves?), or are there connections between phylogeny and specific phenotypes (e.g., does genus X appear only in host Y or in geographic location Z?).

Apart from putting forth novel hypotheses, dbBact makes possible the detection of sources of bacterial groups. We recommend querying dbBact as a first step in any microbiome analysis (e.g., using the interactive heatmap of the dbBact-Calour module). Identifying relevant bacterial groups and their dbBact *annotations* fosters an initial understanding of biological processes, supporting better downstream analysis. dbBact also enables associating bacteria with a “candidate reagent contaminant” *annotation*. We have encountered numerous cases where examining bacteria in a study detected contaminations, e.g., bacteria having mostly “mus musculus” *annotations*, although samples were of human origin. Removing these sequences prior to downstream analysis can remove biases and increase the statistical power of the analysis.

The current coverage of dbBact is high in a large number of habitats, but many other habitats are still poorly covered (e.g., Figure 3e). Therefore, *terms* appearing in a small number of *annotations* may lead to dubious conclusions. For example, dbBact contains a single *experiment* originating from a scrubland environment. This *experiment* profiled the leaf microbiome of ivy plants, hence, querying a set of ivy leaf-related bacteria may result in the enrichment of both “ivy” and “scrubland” *terms*. Therefore, to avoid incorrect conclusions, users are advised to further examine the set of *experiments* associated with each enriched *term*. As the number and diversity of *experiments* in dbBact increase, such spurious *terms* are expected to be suppressed. Tens of microbiome studies are published weekly, but the dbBact team can process only a fraction. We expect the microbiome community to contribute to dbBact and help increase the number and diversity of uploaded studies.

dbBact may also be used in shotgun metagenomics studies. Whenever 16S rRNA sequences are inferred from shotgun data they may be submitted as queries or uploaded to dbBact. The “linking” mechanism for harmonizing studies from different variable regions enables shotgun and amplicon studies to be integrated into one coherent knowledge base. Similarly, studies using long read technologies (or synthetic long reads) also provide full-length 16S rRNA sequences, and thus can be integrated into dbBact in the same manner.

In sum, dbBact introduces a new “layer” of data analysis in microbiome studies. We believe that the scope and ontology-based structure of dbBact provides new means for studying core factors affecting bacterial communities, possibly answering questions that could not have otherwise been asked.

## Supporting information

Supplemental Results and Methods

## Acknowledgements

We wish to thank Tzipi Brown and Rotem Hadar from the Sheba Microbiome Center, Zhenjiang (Zech) Xu, Jon Sanders, Qiyun Zhu, Tomasz Kosciolek, Stefan Janssen and Jeremiah Minich for fruitful discussions, feedback, and suggestions. N.S. is funded by the Ministry of Science, Technology & Space, Israel (Grant 3-16033).

## Methods

As dbBact is constantly growing in scope, and to facilitate reproducibility, the dbBact infrastructure described in this section and all analyses presented in the paper carried out using dbBact release 2022.07.01, available for download as part of the weekly snapshots at https://dbbact.org/download.

### Implementation

#### Database

The dbBact database is stored as a SQL relational database (PostgreSQL 9.5.10). The database schema and detailed table descriptions are provided in Supplementary File 1 and Figures S5-6.

#### Ontologies

Table S1 presents ontologies available in dbBact release 2022.07.01. dbBact supports the addition of ontologies to allow more accurate annotations. When users provide *terms* that do not appear in any of these ontologies, a new *term* is automatically added to the generic dbBact ontology.

#### ASV sequences

##### Primers and trimming

dbBact uses exact prefix search for *sequence* identification, and therefore all *sequences* in dbBact are primer trimmed and originate from one of the supported 16S rRNA forward primers. For dbBact release 2022.07.01, the supported forward primers are V1-27F (AGAGTTTGATCMTGGCTCAGxxx), V3-341F (CCTACGGGNGGCWGCAGxxx), V4-515F (GTGCCAGCMGCCGCGGTAAxxx), where “xxx” denotes the beginning of the ASV sequence stored in dbBact. Although additional primers can be added to dbBact, the vast majority of 16S rRNA studies uses one of the three primers described. The minimum length of *sequences* uploaded to dbBact is 100bp. Upon upload, *sequences* are stored at their full length rather than being truncated to a fixed length. When submitting a query *sequence*, exact sequence matches are searched using *length=min(query_sequence_length, database_sequence_length)*.

##### Taxonomy assignment

A python script runs daily to add taxonomy assignments to uploaded dbBact ASV *sequences* using RDP version 2.12 (69). Although taxonomy is not used in dbBact for analysis, it may be used for querying dbBact (e.g., retrieving *annotations* associated with bacteria of the genus *Streptococcus*).

##### Inter-region querying

dbBact supports the harmonization of microbiome studies performed using different protocols by inter-region linking. When submitting a *sequence*, dbBact uses the SILVA database of full length 16S rRNA genes (SILVA version 132, (34)) to identify *sequences* whose “footprint” in other variable regions matches the query. First, the SILVA sequences containing the query *sequence* are detected. Second, all dbBact *sequences* that match these SILVA sequences, in any region, are retrieved (i.e., Query(S)=*{T: T ∈ dbBact, ∃R ∈ SILVA so that ss(S, R) and ss(T, R)}* where *ss* stands for “subsequence”). Querying is performed using the “wholeSeqIDsTable” table in the dbBact implementation. To enable fast queries, a daily script is run on new dbBact *sequences*, linking all *sequences* sharing a SILVA sequence. Such linking is performed only when querying dbBact, therefore new versions of SILVA or other full length 16S rRNA databases may be seamlessly applied. Currently, dbBact supports linking the V1, V3, and V4 forward primer reads; additional primers may be incorporated if needed.

##### Queries of different sequence length

*Sequences* uploaded to dbBact may vary in length depending on the sequencing platform and sequenced region. When adding new *annotations*, dbBact stores the full-length sequence of each ASV. For example, when two *experiments* provide information about the same bacterium using 150bp and 200bp reads, respectively, dbBact stores these *sequences* as separate entries and links each *annotation* to the corresponding *sequence*. Yet, when submitting a query using either *sequence*, dbBact retrieves *annotations* using exact match on the shortest common sequence, hence also retrieving *annotations* related to the other *sequence*.

### dbBact interfaces

#### REST-API server

The dbBact REST-API server (http://api.dbbact.org) is implemented in Python 3.6, using Flask version 0.12/Gunicorn v19.9 to handle web queries, and psycopg2 version 2.7.1 for handling Postgres data queries. Full API documentation is available at http://api.dbbact.org/docs. Examples using the REST-API for querying are available at: https://github.com/amnona/dbbact-examples. The REST-API enables access to all dbBact functions. Querying dbBact or adding anonymous *annotations* does not require registration. Registration by username/password enables editing *annotations* submitted by the same user.

#### dbBact website

The dbBact website (http://dbbact.org) enables dbBact *annotation* retrieval based on ASVs, taxonomy, or ontology *terms*. Additionally, the website provides word-cloud generation and *term* enrichment analysis. The source code for the website as well as deployment instructions are available on the dbBact-website github page (https://github.com/amnona/dbbact-website).

#### dbBact-Calour interface

dbBact is integrated into the Calour microbiome analysis program (https://github.com/biocore/calour), using the dbBact-Calour module (https://github.com/amnona/dbbact-calour). Using this interface, users can both query dbBact regarding bacterial sequences, and add new *annotations*. The dbBact-Calour module provides dbBact *annotation* retrieval from the interactive Calour heatmap display, showing all *annotations* associated with the selected *sequence*. Additionally, the module enables GUI-based creation of new dbBact *annotations* for selected *sequences*, and performs *term* enrichment analysis, *term*-based PCA and word cloud generation. A Jupyter notebook tutorial is available at: http://biocore.github.io/calour/notebooks/microbiome_databases.html.

The module also works with EZCalour, the full GUI version of Calour, (https://github.com/amnona/EZCalour). A tutorial for dbBact enrichment analysis using EZCalour is available at: https://github.com/amnona/EZCalour/blob/master/using-ezcalour.pdf.

#### Qiime2 plugin

The q2-dbBact plugin (https://github.com/amnona/q2-dbbact) enables dbBact *annotation*-based analysis using the qiime2 framework (35). The interface provides *term* enrichment analysis for the output of various qiime2 differential abundance plugins (ANCOM (70), Songbird (71), ALDEx2 (72), DACOMP (73), or a rank-mean method). Additionally, the plugin supports dbBact *term* word cloud and interactive heatmap generation.

### Data availability

Table 1 details the locations of different dbBact sites

**Table 1.** Locations of dbBact sites

|  | Location |
| --- | --- |
| dbBact website | http://dbbact.org |
| dbBact website source code | https://github.com/amnona/dbbact-website |
| dbBact REST-API server | api.dbbact.org |
| Documentation for the REST-API | api.dbbact.org/docs |
| Examples for using the REST-API interface | https://github.com/amnona/dbbact-examples |
| Source code for the REST-API server | https://github.com/amnona/dbbact-server |
| Jupyter notebooks for figures presented in the paper | https://github.com/amnona/dbbact-paper |
| Weekly dump of the complete dbBact database<br>(excluding user details) | https://dbbact.org/download |

### Standard analysis: Default dbBact preprocessing of an *experiment*

Although dbBact is a wiki-style knowledge base, the vast majority of *annotations* in dbBact release 2022.07.01 was added by the dbBact team. Studies were selected from published microbiome papers, and *annotations* were added following the re-processing of the experimental data, using a “standard” manual analysis pipeline as follows:

The raw data of each scientific paper (i.e., per-sample FASTA files and corresponding metadata) were downloaded using the provided accession (e.g., by SRA/ENA accession or Qiita (21) study ID). When data or metadata were not available, the authors were contacted and provided the missing data directly. When primer sequences were part of the reads, they were removed using a custom script (https://github.com/amnona/GetData). Subsequently, the Deblur pipeline (15) was applied to the reads of each sample (Deblur script version 1.1.0, using default parameters, https://github.com/biocore/deblur), resulting in a denoised biom table.

This biom table, together with the per-sample metadata, were manually re-analyzed using Calour (36), to add *annotations* capturing biological conclusions arising from the study. Three types of predicates were sought:

(i) “DIFFERENTIAL:” To detect sets of *sequences* associated with relevant conditions (e.g., sick vs. healthy), *sequences* significantly enriched between two conditions were identified using a non-parametric permutation based rank-mean test, followed by multiple hypothesis correction using dsFDR (74) (usually set to 0.1). The test was performed by the calour.diff_abundance() function. Correlations with continuous metadata fields (e.g., BMI) were detected with a permutation-based Spearman test with dsFDR correction using calour.correlation(). In both cases, the set of *sequences* higher or lower in one condition than in the other were then annotated as “DIFFERENTIAL,” i.e., “HIGHER IN” condition 1 and “LOWER IN” condition 2.

(ii) “COMMON”/“DOMINANT:” For each study, *sequences* present in more than half of the relevant samples were *annotated* as “COMMON.” “DOMINANT” *sequences* were identified as *sequences* whose mean frequency in the relevant samples was higher than 0.01. In studies containing samples from multiple sources (i.e., fecal and saliva samples, or samples from individuals from several countries or disease vs. healthy), a “COMMON”/“DOMINANT” *annotation* was added separately to each source subset.

(iii) “CONTAMINANT:” When a study contained a set of negative control (blank) samples, ASVs showing higher frequency in these controls than in the non-blank samples were manually annotated as a possible “CONTAMINANT.”

Examples of the different predicates appear in Table S3.

#### Remark

Using the abovementioned pipeline is not a prerequisite for adding new *annotations*, and any denoising method followed by statistical analyses can be applied by users contributing to dbBact.

### Statistical analysis in dbBact

#### Word cloud generation

*Calculating a term’s* F1 *score*: Given a set of query *sequences S*, each dbBact *term t* is assigned an F1 score, corresponding to the harmonic mean of precision and recall. For each sequence *s* in *S*, we calculate the the fraction of *s*’s *annotations* that contain the *term t*. The average of these values across *S* provides the precision of *t* on *S*. Similarly, for a given *sequence s* and a *term t*, recall is calculated as the fraction of *t*’s *annotations* that contain *s*. To suppress terms that appear in a small number of *experiments*, the total number of *t*’s *annotations* (i.e., the denominator) is artificially increased by 1. The average of these values across *S* provides the recall of *t* on *S*.

*Displaying a word cloud*. The word cloud size of each *term* is proportional to its F1 score. If the term appears in “LOWER IN” annotations, its color is orange, otherwise it is blue. The brightness of each *term* represents the number of *experiments* containing the *term*, indicating the reliability of the term (white for a single *experiment*; ranging to dark blue/orange for >=10 *experiments*).

Word clouds were generated with the dbBact-Calour module for Calour (https://github.com/amnona/dbbact-calour) using the draw_wordcloud() function.

#### Term enrichment analysis

Given two sets of *sequences, S1* and *S2*, we search for *terms* significantly enriched in either group using the following steps:

*(a) Calculating an annotation-based score per sequence and term*. Each *annotation a* in dbBact is assigned a “weight” *w(a)* according to its predicate. The predicates “COMMON,” “CONTAMINATION,” and “DIFFERENTIAL” are assigned a weight of 1, and the predicates “DOMINANT” and “OTHER” are assigned a weight of 2 and 0.5, respectively. These weights are applied to calculate a score *Score(s, t)* for each *term t* in dbBact and a *sequence s* in either *S1* or *S2*. The score sums the weights of all *annotations* involving *s* and *t*. When *t* appears as “LOWER IN” in the predicate “DIFFERENTIAL,” a new *term* “not *t*” is created and assigned a weight of 1.

*(b) Calculating effect size of a term*. For a *term t, Score(s, t)* is calculated for all *sequences* in *S1* and *S2*, and the effect size of *t* is defined as

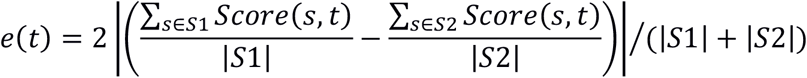

where |*S*| corresponds to the number of *sequences* in the set *S*.

*(c) Finding significant terms*. Each *term* is assigned a p-value by comparing its scores over 1000 random permutations of the combined *S*1 and *S*2 *sequences* to sets of size |*S*1| and |*S*2|. Subsequently, a dsFDR multiple hypothesis correction (with a threshold of 0.1) is applied to detect significant *terms*.

*(d) Calculating the significance of a term across experiments*. Until this stage, we measured the enrichment of a *term* based on all dbBact *experiments* combined. To estimate whether such significance occurs across multiple *experiments*, or whether it is driven by a single or a few *experiments*, steps (a)-(c) were also repeated using each individual *experiment* that contains *t*. The total number of *experiments* containing each *term* and the fraction in which the *term* was significant appear in each figure.

The abovementioned analysis is performed using the dbbact-calour module enrichment() function.

#### Venn diagrams

Given two sets of sequences, *S*1 and *S*2, and a *term t*, we plot a Venn diagram indicating the number of *sequences* associated with *t* across all dbBact *annotations*, and the overlap of these *sequences* with *S*1 and *S*2.

Venn diagrams were generated using the dbbact-calour module plot_term_venn_all() function.

#### dbBact *term-*based principal component analysis

Each sample *x* is represented by a vector of *sequence* frequencies (abbreviated “sf”), *x(S) = (sf*1, *sf*_2_, *…, sf*_*n*_) across the *n sequences* that appear in all study samples (*sf* equals zero in case a *sequence* does not appear in a specific sample). We then transform *x(S)* into a *term*-based representation of *x*, i.e., *x(T) = (ts*1, *ts*_*2*_, *…, ts*_*m′*_), where each *ts* is a *term*-score described below, calculated across all *m* dbBact *terms*. The *ts* score for the *term t* is given by 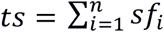 *precision(i, t)*, where *precision(i, t)* is the fraction of *annotations* associated with *sequence i* that also contain the *term t*. Finally, once *x(T)* is calculated over all samples in a study, we perform principal component analysis of this space. Each principal axis is defined by its weights, where the highest weights (in absolute value) are used for providing biological meaning. The abovementioned analysis is performed using the dbbact-calour module plot_term_pcoa() function.

### Processing of datasets

All datasets discussed in the paper were processed using the following pipeline: raw reads were downloaded and denoised using the Deblur pipeline (15) with default parameters; the resulting denoised biom table was loaded into Calour (36), and differentially abundant bacteria were identified using a permutation-based non-parametric rank mean test with dsFDR multiple hypothesis correction (74) set to 0.1 (using the calour diff_abundance() method). In the case of the American Gut Project dataset, multiple samples originating from the same individual were aggregated to a single sample using mean frequency for each ASV. The groups of high and low fruit consumption were controlled for confounders by stratifying samples in both groups based on the AGP metadata fields: age category (“AGE_CAT”), sex and BMI category (“BMI_CAT”), and randomly dropping samples to equalize the number of samples from each stratum prior to differential abundance testing. dbBact *term* word clouds were generated by applying the above-described word cloud approach using the dbbact-calour module draw_wordcloud() method on all sequences present in at least 30% of the samples. *Term* enrichment was performed using the above-described term enrichment approach using the dbbact-calour module enrichment() method with default parameters (dsFDR=0.1). dbBact term PCAs were generated using the dbbact-calour module plot_term_pcoa() function. Accession numbers for each dataset used are available in Supplementary File 2. Jupyter notebooks used for the creation of each figure are available at https://github.com/amnona/dbbact-paper.

