## Supplemental Results and Methods for "Achieving pan-microbiome biological insights via the dbBact knowledge base"

### Supplementary Results

#### Distribution of dbBact experiments across different ontologies

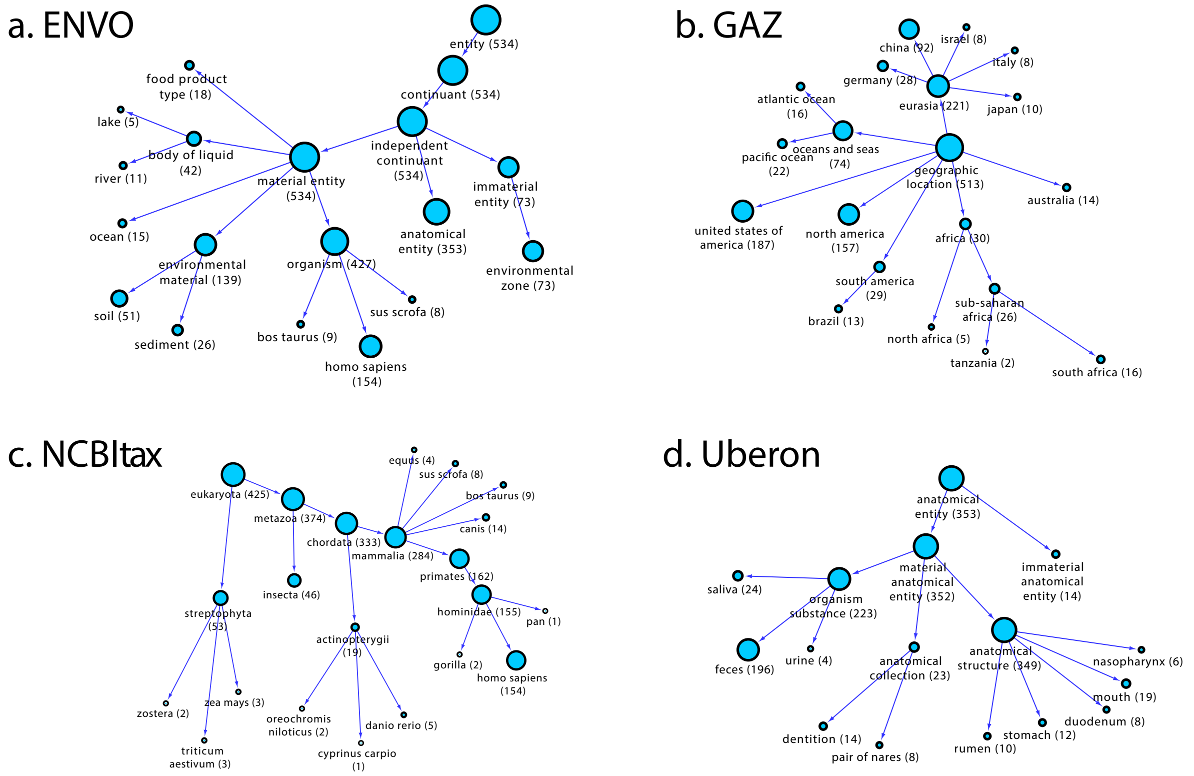


**Figure S1: Distribution of dbBact experiments by different ontologies**. The number of *experiments* containing representative terms from: **a.** ENVO, covering environment types (1,2). **b.** GAZ, specifying geographic locations (3). **c.** NCBI taxonomy of species (4). **d.** Uberon, covering anatomical entities (5). Numbers in parenthesis and circle sizes denote the number of unique dbBact *experiments* containing these *terms*.

#### Querying *sequences* from one region and retrieval across all regions

When submitting queries from regions V1-V2, V3-V4 or V4, *sequences* are matched with full-length 16S rRNA sequences based on the SILVA database, then "linked" to the corresponding *sequences* in the other regions (see Methods section for implementation details). The latter corresponding sequences are automatically submitted as queries to dbBact, therefore retrieval is based on *annotations* across all regions. For example, when submitting a query *sequence* from the V1-V2 region, dbBact also "transforms" it into its matching *sequences* in V4 and in V3-V4 and retrieves also *annotations* that were originally submitted using these regions.

To exemplify this process, we submitted queries of V3-V4 *sequences* from soil samples (6), V3-V4 *sequences* from mouse and pig feces (7,8), and V1-V2 *sequences* from human feces (9). Figure S2a displays dbBact word clouds for each sample when retrieval was restricted to V4 linked *sequences*, showing accurate detection of the origin of the *sequences* based only on the linked *annotations*. Figure S2b displays a similar example for an enrichment query of 55 and 332 V1-V2 *sequences* that are higher in short bowel syndrome (SBS) patients and healthy controls, respectively. Although dbBact retrieval was restricted to V4 linked *sequences*, the bar plot identifies an enrichment of general dysbiosis bacteria (oral and diarrhea associated) in the SBS group, as well as an enrichment of non-dysbiosis bacteria ("LOWER IN ulcerative colitis") in the control group. This indicates the SILVA-mediated sequence linking can assist in identifying subtle biological signals even for primer regions lacking direct experiments for these signals.

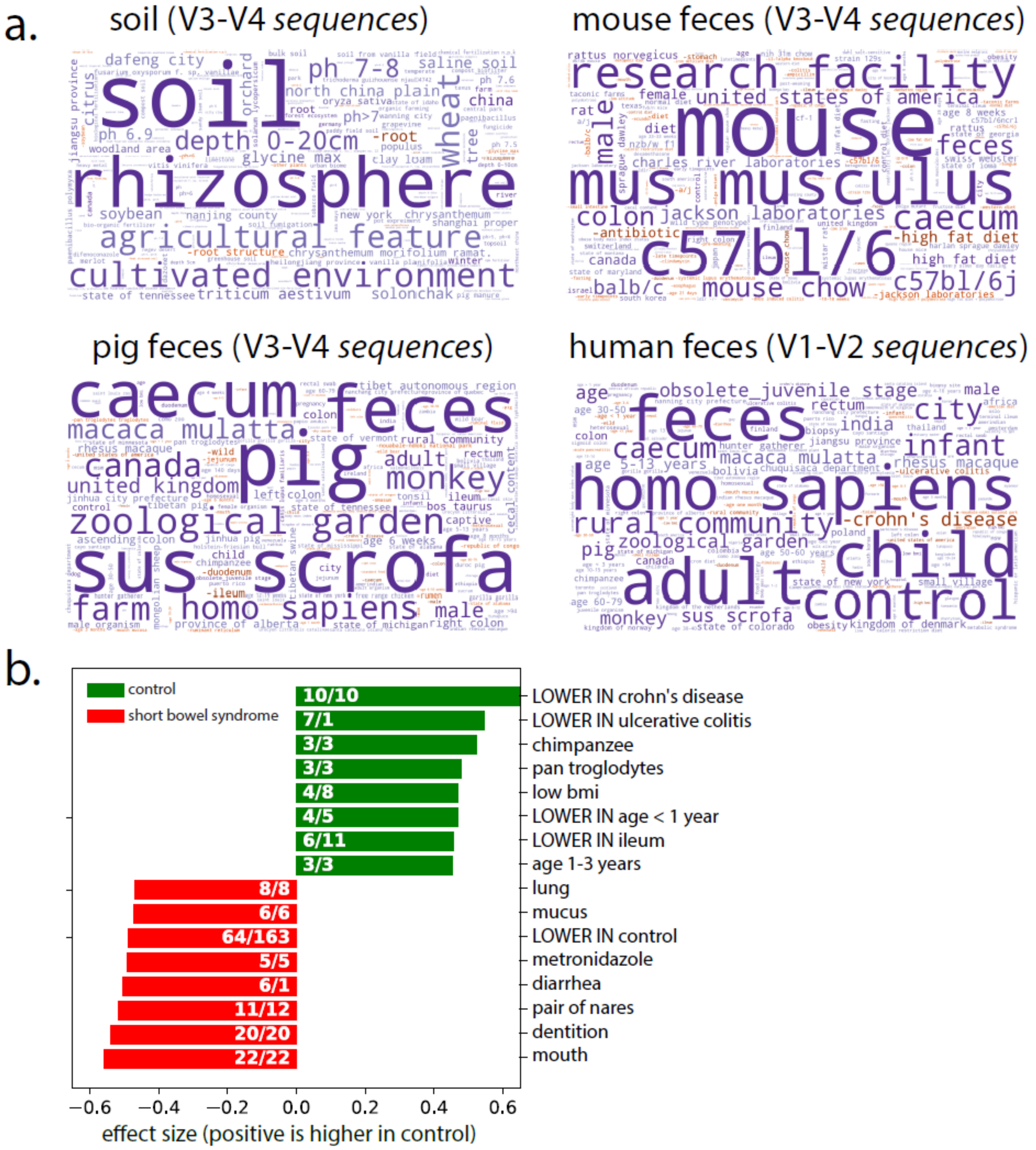


**Figure S2: Querying dbBact using V3-V4 or V1-V2 16S rRNA gene regions and retrieving *annotations* from V4**. **a.** Reads originating from soil (6) (V3-V4 region), mouse feces (8) (V3-V4 region), pig feces (7) (V3-V4 region), and human feces (9) (V1-V2 region) were used to query dbBact, while retrieval was restricted to their linked V4 *sequences*. Word clouds based on V4 *annotations* are shown for each sample. **b.** Top dbBact *terms* significantly enriched between SBS patients and healthy controls using only V4 region *annotations*, while the two-group enrichment query was based on V1-V2 reads. Results indicate general dysbiosis of SBS bacteria (similar to Crohns' disease, ulcerative colitis, and pancreatitis), as well as enrichment of mouth bacteria in SBS patients.

#### Integrating dbBact into microbiome analysis pipelines allows generating novel biological hypotheses: Detailed examples

The following sections present a detailed analysis of examples mentioned in Figure 6a.

Jupyter notebooks for recreating all analyses and figures, as well as the data used, are available at: https://github.com/amnona/dbbact-paper

#### Oral microbiome of wild sea otters resembles the oral microbiome of dogs and cats

Dudek et al. (10) collected oral samples from 158 wild southern sea otters living off the coast of central California. Figure S3a displays the *terms* word cloud of all sea otter oral samples indicating shared *sequences* with dogs and cats (e.g., "beagle dog," "canis lupus familiaris," "felis catus"). To validate and further understand this observation, we plotted the fraction of sea otter oral *sequences* shared with oral or non-oral *terms* corresponding to dogs, cats, humans, and fish (Figure S3b). For example, dbBact contains 19 *annotations* of dogs' oral microbiome (leftmost column in Figure S3b). Comparing the *sequences* corresponding to each of these *annotations* with the set of 125 sea otter oral *sequences* reveals an overlap of up to 30% (median 10%) across *annotations*. A significantly lower overlap exists when comparing the 125 sea otter *sequences* with *sequences* corresponding to non-oral dog *annotations*. The same phenomenon occurs for cats' oral annotations. By contrast, oral and non-oral *annotations* from humans and fish show a low overlap with the 125 sea otter oral *sequences* (Mann-Whitney p-values <1E-8 for all comparisons). These results indicate that dogs' and cats' oral bacteria are the closest to sea otter oral samples, possibly because of a common local environment and diet.

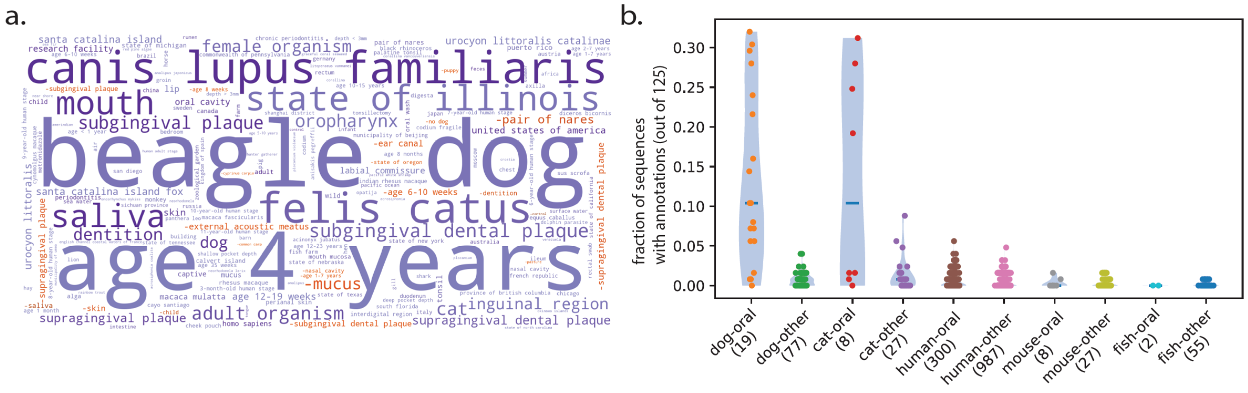


**Figure S3: Sea otter oral microbiome resembles the oral microbiome of dogs and cats.** **a.** dbBact *term* word cloud of prevalent *sequences* (present in >0.3 of samples) of wild sea otter oral samples. **b.** Fraction of the 125 sea otter *sequences* (present in >10% of samples) that also appear in *annotations* containing specific dbBact *terms*. Each point represents a single dbBact *annotation*. To collect oral-related *annotations* we used the *terms* "mouth," "saliva," "dentition," or "oropharynx," together with the host *term* (i.e., "dog," "cat," "homo sapiens," or "fish"). Non-oral *annotations* were taken as all other *terms* for each host. Numbers in parenthesis correspond to the number of *annotations*.

#### Himalayan vulture fecal microbiome resembles that of the California condor

Wang et al. (11) collected 28 fecal samples from wild Himalayan vultures in Qinghai province, China. Figure S4a shows the most significant dbBact *terms* associated with the 36 *sequences* present in at least 30% of Himalayan vulture samples. The two dbBact *terms* having the highest associated $F_1$ scores are "feces" and "gymnogyps californianus" (California condor). About 60% of the Himalayan vulture *sequences* are associated with "California condor," and over 80% are associated with "feces" (Figure S4b). dbBact California condor *annotations* are derived from the Jacobs et al. *experiment* (12), which contains fecal and cloacal samples of captive California condor collected in Idaho, USA. Thus, dbBact indicates a common bacterial core for these two carrion-eating vultures from different continents.

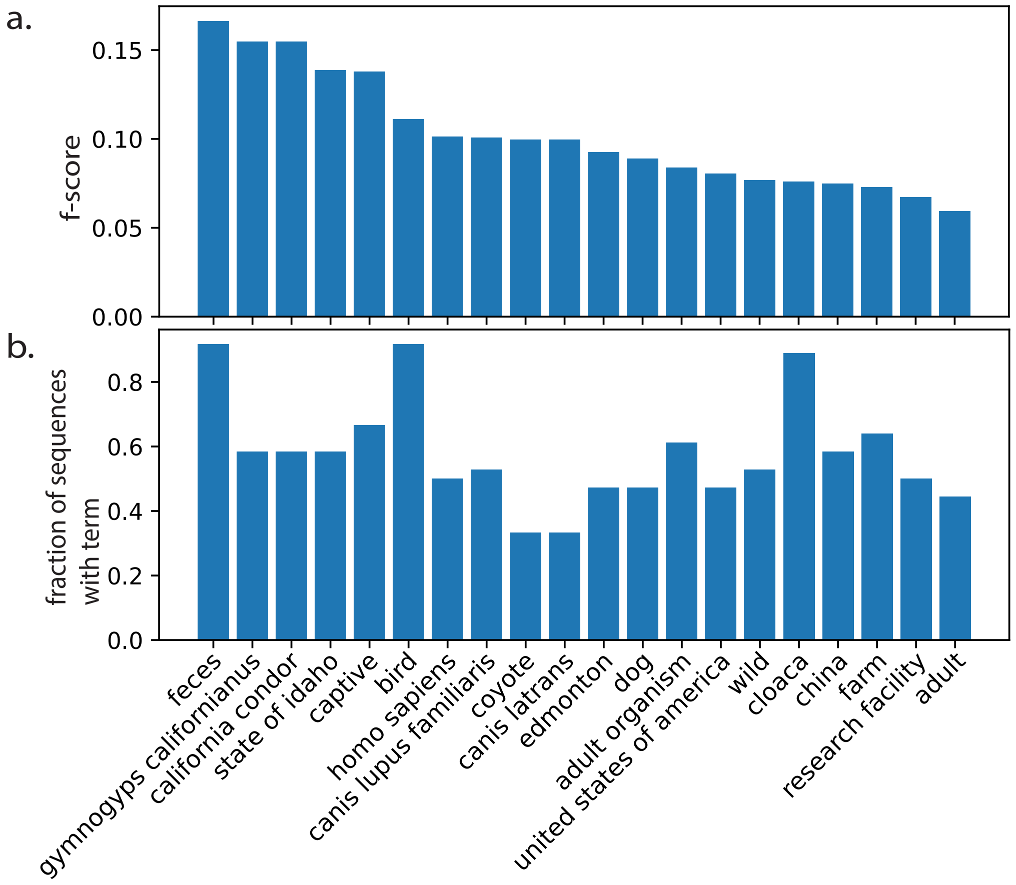


**Figure S4: Himalayan vulture microbiome resembles that of the California condor. a.** dbBact *terms* with the highest $F_1$ scores. **b.** The fraction of Himalayan vulture *sequences* (present in >30% of samples) associated with each of the *terms* shown in panel a.

##### Colitis in horses increases the abundance of human-related bacteria in the horse gut

Arnold et al. (13) collected fecal samples from 80 healthy horses and 26 horses suffering from colitis due to *Salmonella* or antibiotics use. Standard analysis identified 2,441 *sequences* higher in healthy horses (S-Normal) and 399 *sequences* higher in horses with colitis (S-Colitis). dbBact *term* enrichment indicates that S-Normal is enriched in horse and other species of the same genus ("equus caballus," "equus hemionus), whereas S-Colitis is enriched in human related terms ("homo sapiens," "adult," "child") (Figure S5a). For example, we compared S-Colitis *sequences* with all 6,037 dbBact *sequences* having an *annotation* "child." 26% (102/399) of S-Colitis *sequences* were independently assigned a "child" *annotation* across dbBact, as opposed to only 1.7% (42/2,441) of S-Normal *sequences* (Figure S5b). This enrichment in human-related *terms* may indicate a decrease in host-specific *sequences* and an increase in non-specific opportunistic *sequences*, possibly arising from horses' exposure to human bacteria following colitis.

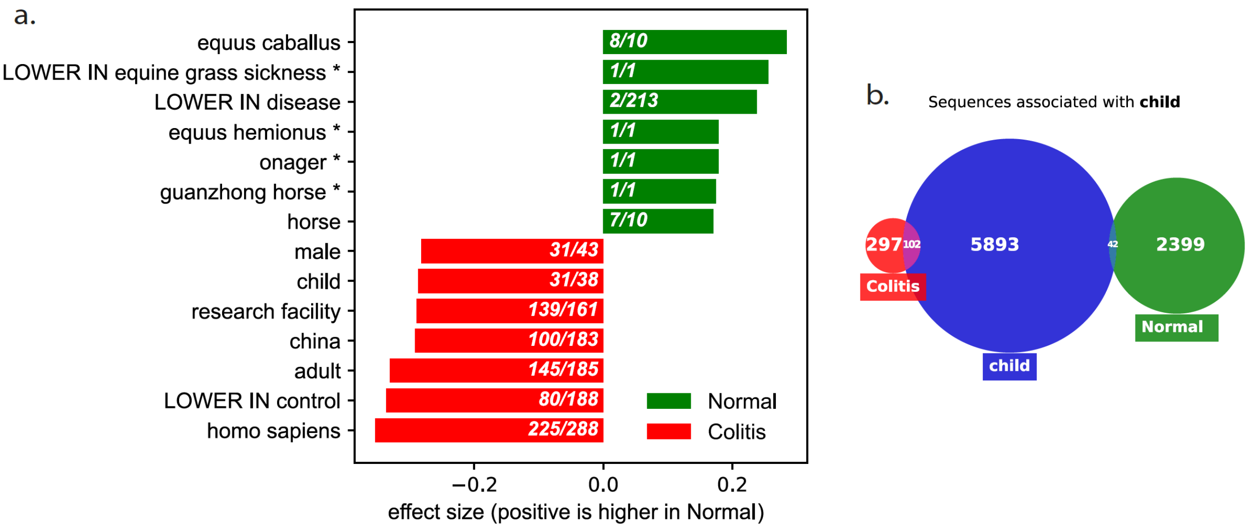


**Figure S5: Bacteria of horses with colitis are more human related. a.** Bar plot of the top 10 enriched dbBact *terms* comparing the *sequences* higher in S-Normal (green) and S-Normal (red). **b.** Venn diagram showing the number of *sequences* associated with at least one dbBact *annotation* containing the *term* "child." Green and red circles are the *sequences* of the S- Normal and S-Colitis, respectively. The blue circle corresponds to all dbBact *sequences* associated with the *term* "child."

#### Human health-related bacteria are associated with primates, whereas disease-related bacteria are associated with homeothermic hosts

In a recent meta-analysis, Abbas-Egbariya et al. (14) identified a set of bacteria that displayed the same disease-dependent behavior across multiple studies and diseases. They identified a set of 34 disease-related bacteria whose frequency consistently increased in different diseases, and a set of 97 bacteria whose frequency decreased in the various disease types, which are referred to as health-related. We used dbBact to identify enriched *terms* in these two groups (Figure S6a). Disease-related bacteria are enriched in "Crohn's disease" and "LOWER in controls," whereas health-related bacteria are enriched in "LOWER-in-disease" *terms* (e.g., "LOWER in Crohn's disease," "lower in pancreatitis," "lower in ulcerative colitis").

Disease-related bacteria are also enriched in young age *terms* (e.g., "infant," "6-month old human stage," "under-1-year human stage"), indicating a possible state of dysbiosis associated with immature microbiome populations. Conversely, health-related bacteria are also enriched in rural lifestyle associated *terms* ("small village," "rural community") and caloric restriction diet ("cron diet").

To gain further insight into the factors determining this cross-disease bacterial response, we repeated *term* enrichment, excluding human disease-associated *annotations* (i.e., analysis was limited to *annotations* that do not contain the *term* "homo sapiens") (Figure S6b). Health-related bacteria are also observed in multiple primate *annotations*, e.g., chimpanzee (69 out of 97 health-related bacteria were observed in chimpanzees compared to only 1 out of the 34 disease-related bacteria; Figure S6c). By contrast, disease-related bacteria are enriched in mouse, horse, rat, and chicken *annotations* (Figure S6d). These results indicate that the bacteria disappearing in various diseases are more specifically adapted to humans, whereas the ones appearing in disease are more generalist homeothermic host-associated bacteria, and therefore probably less specifically adapted to their human host.

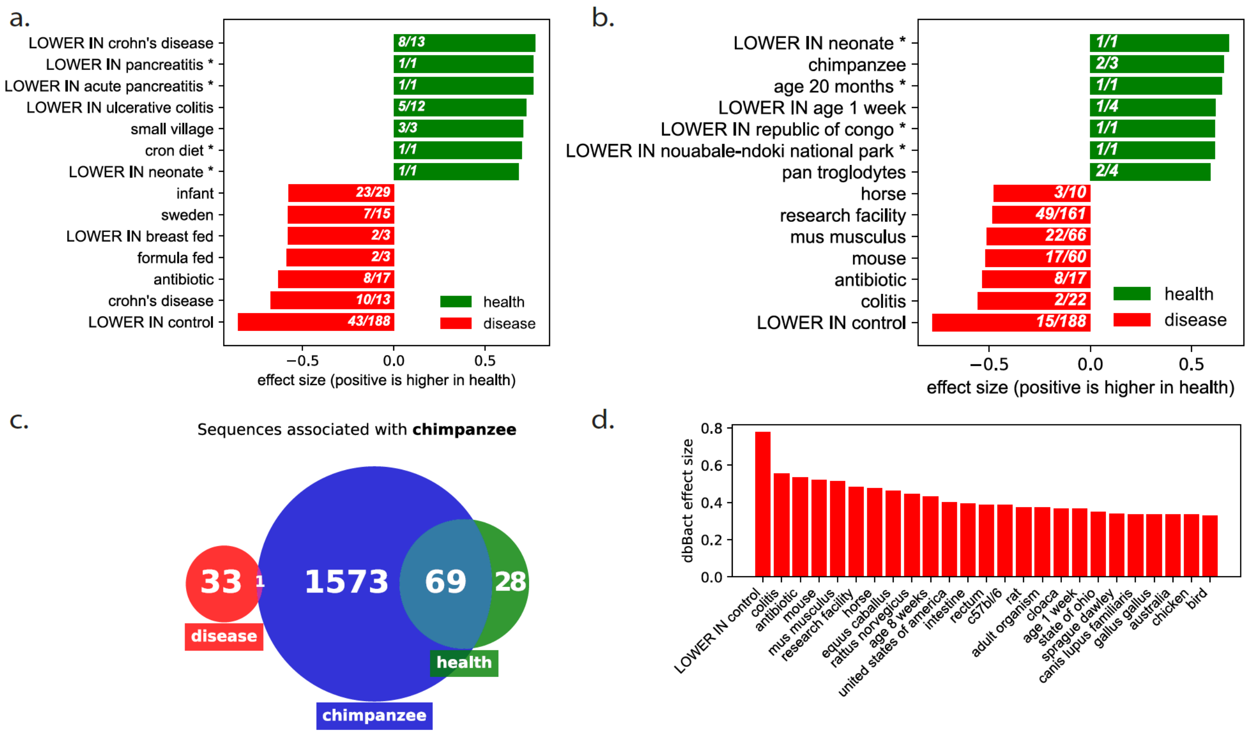


**Figure S6: Health-related bacteria are associated with primates, whereas disease-related bacteria are associated with other homeothermic hosts**. **a.** Bar plot of the top 10 enriched dbBact *terms* comparing the health-related sequences (green) and disease-related *sequences* (red). **b.** The same as a. but restricting enrichment analysis to non-human *annotations*, i.e., those that do not contain the *term* "homo sapiens." **c.** Venn diagram of dbBact *annotations* related to the *term* "chimpanzee." Green and red circles indicate the number of *sequences* associated with the *term* in the health- and disease-related bacteria, respectively; the blue circle indicates the number of "chimpanzee" *sequences* across dbBact. **d.** The top 25 *terms* of the disease group in b.

#### High fruit consumption is associated with monkey- and rural- associated bacteria

Samples from the American Gut project (15) were filtered based on levels of fruit consumption (higher/lower than three fruits per week). Samples were selected so that the distributions of age, BMI, and sex were similar, resulting in 1,071 samples in either low or high fruit consumption groups (see Methods section "Processing of datasets" for more details). Standard analysis yielded 45 *sequences* higher in high fruit consumption and 41 *sequences* higher in low fruit consumption, referred to as S-high and S-low, respectively. dbBact enrichment analysis (Figure S7a) shows that S-low *sequences* were associated with diets in industrialized regions (e.g., "LOWER IN rural community"), and S-high was enriched with the *terms* "hunter gatherer," "monkey," and rural environment-related *terms*. For example, the *term* "monkey," appearing 5,292 times in dbBact, is associated with 86% (39/45) of S-high *sequences*, compared to 12% (5/41) of S-low (Figure S7b). This may indicate that the majority of bacteria positively affected by a high fruit diet are universal responders to fruit consumption, and thus also appear in monkeys. Therefore, some of the differences in microbial communities between humans and monkeys are due to diet rather than host-microbe co-evolution.

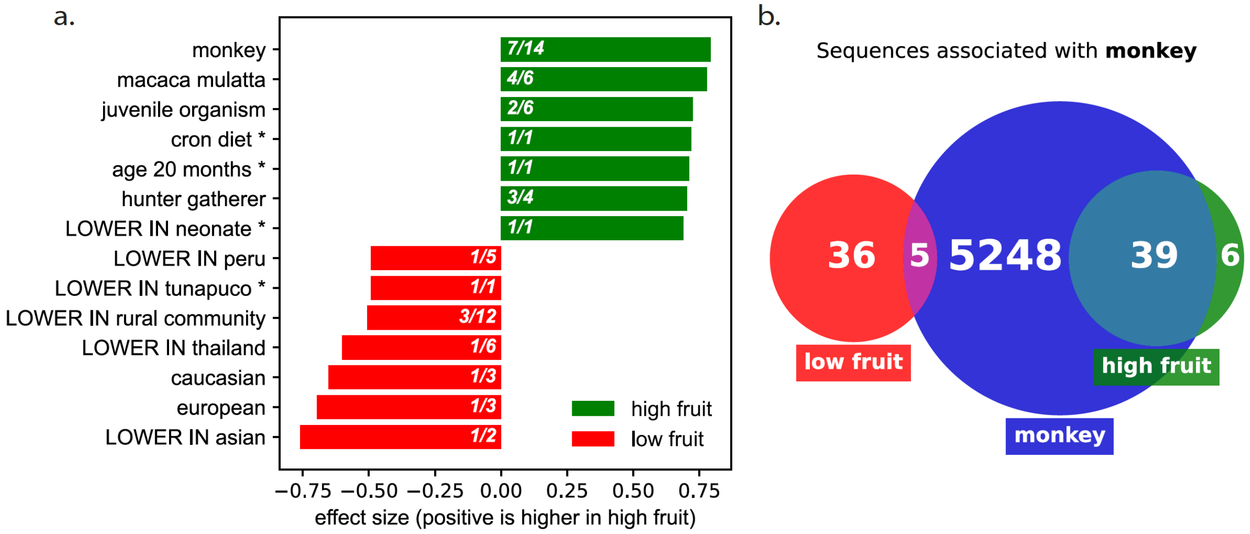


**Figure S7: High fruit consumption is associated with monkey- and rural-associated bacteria**. **a.** Bar plot of enriched *terms* in high/low fruit consumption groups. **b.** Venn diagram for the representative *term* "monkey," displaying the number of dbBact *sequences* associated with the *term* (blue circle), and their overlap with *sequences* associated with that *term* in either group (red or green circles). The numbers indicate the number of *sequences* in each part of the Venn diagrams.

#### Enrichment of oral related bacteria in fecal IgA-positive compared to IgA-negative fraction

Scheithauer et al. (16) collected fecal samples from individuals with obesity. Amplicon sequencing was applied to the IgA positive and IgA negative fractions of the fecal microbiome following anti-human-IgA FACS. Using standard analysis, we looked for differentially abundant bacteria in the IgA positive vs. IgA negative fractions (i.e., comparing IgA positive and negative samples from each subject). An FDR threshold of 0.5 was used to obtain a large set of features to enable robust dbBact *term* enrichment, resulting in 70 and 137 *sequences* significantly higher in the IgA negative and positive fractions, respectively.

dbBact *term* enrichment (Figure S8a) shows that the IgA negative fraction is enriched in IgA negative bacteria from other *experiments* (e.g., "IgA negative fraction," "only lower in IgA positive fraction"), indicating a conserved signal of IgA binding across studies. In addition, the IgA positive fraction is enriched in oral associated *terms* (e.g., "oral cavity," "oral wash," "lower in dentition," "dentition"). For example, *annotations* of the *term* "oral cavity" contain 1990 *sequences*, out of which 36 are shared with *sequences* of the IgA positive fraction, and none with the IgA negative fraction (Figure S8b). It is reasonable to hypothesize that oral bacteria are more likely to be IgA bound, possibly because of binding in the oral cavity.

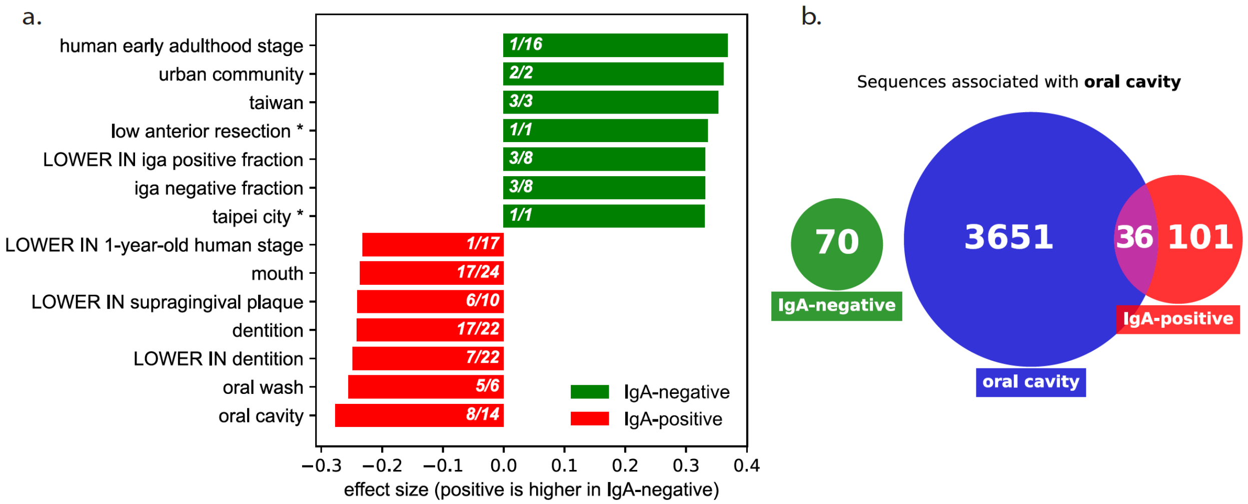


**Figure S8: Enrichment of oral related bacteria in fecal IgA-positive compared with IgA-negative fraction. a.** Bar plot of the top 10 enriched dbBact *terms* comparing the *sequences* higher in IgA negative (green) and IgA positive (red) fecal samples. **b.** Venn diagram showing the number of *sequences* associated with the *term* "oral cavity." Green and red circles are *sequences* higher in the IgA negative and IgA positive fractions, respectively. The blue circle corresponds to all dbBact *sequences* associated with the *term* "oral cavity."

Braces lead to enrichment of dentition-related bacteria in saliva

Willis et al. (17) collected oral rinse samples from age-matched adolescent students with and without braces. We tested for *sequences* significantly higher in participants with/without braces, resulting in sets of 146 and 40 *sequences*, respectively. dbBact *term* enrichment (Figure S9) shows that the no-braces group is enriched in non-dentition *terms* (e.g., “tongue,” “lower in supragingival plaque,” “lower in dentition”), whereas participants with braces show strong enrichment in dentition-related *terms* (e.g., “subgingival dental plaque,” “dentition,” “lower in saliva,” “supragingival plaque”). The latter enrichment may be due to bacterial attachment to the braces.

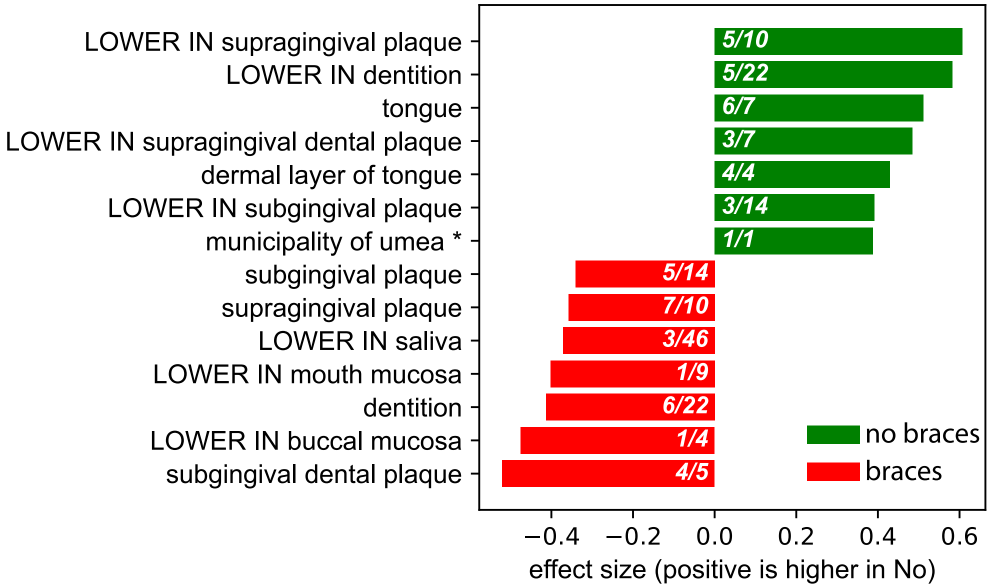


**Figure S9: Braces cause enrichment of dentition-related bacteria.** Bar plot of the top 10 enriched dbBact *terms* comparing the *sequences* higher in children with and without braces (red and green, respectively).

##### Oral rinse bacteria of acute tonsillitis patients are enriched in soft-tissue-associated bacteria, and are depleted of dentition-associated bacteria

Yeoh et al. (18) compared the oral cavity bacteria of 43 acute tonsillitis patients with that of 165 individuals without tonsillitis. Standard analysis comparing the two groups detected 29 *sequences* higher in tonsillitis patients and 59 *sequences* higher in non-tonsillitis controls, referred to as S-tonsillitis and S-non, respectively. dbBact enrichment analysis (Figure S10) shows that S-non *sequences* are enriched in dentition-associated *terms* (e.g., "supragingival plaque," "dentition," "LOWER in saliva," "subgingival plaque"), whereas S-tonsillitis sequences are enriched in soft tissue-associated terms (e.g., "tongue," "LOWER IN supragingival plaque," "LOWER IN dentition"). Oral rinse represents sampling of bacteria from multiple oral cavity sources, but typically tonsil-associated bacteria are not expected to be sampled in saliva. Therefore, the increase in soft-tissue-associated bacteria (and the decrease in dentition-associated bacteria) in tonsillitis patients could be due to increased amounts and/or increased shedding of non-tonsil soft-tissue-associated bacteria.

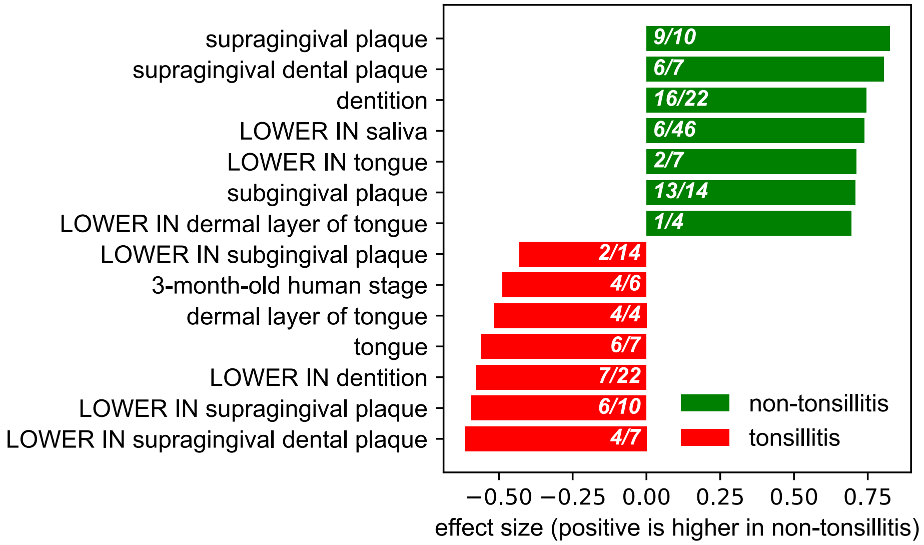


**Figure S10: Oral rinse bacteria of acute tonsillitis patients are enriched in soft-tissue-associated bacteria and depleted of dentition-associated bacteria**. Bar plot of enriched dbBact *terms* comparing the *sequences* higher in tonsillitis (red) and non-tonsillitis (green).

#### Acute pancreatitis and Crohn’s disease share gut bacteria

Fecal samples of 135 acute pancreatitis patients (seven days from the onset of symptoms) and of 35 healthy controls were analyzed (19). Standard analysis detected 39 *sequences* higher in pancreatitis patients and 296 *sequences* higher in healthy controls, referred to as S-panc and S-healthy, respectively. dbBact enrichment analysis shows that S-healthy *sequences* were enriched in rural and health-associated *terms* such as “LOWER in crohn’s disease” and “rural community,” whereas enriched *terms* in S-panc *sequences* included disease-associated *terms* such as “diarrhea,” “crohn’s disease,” and “LOWER in control,” as well as oral *terms* such as “saliva” and “dentition” (Figure S11a). For example, 36% (14/39) of the S-panc *sequences* were independently annotated as “crohn’s disease”-associated *sequences* across dbBact, compared to only 14% (41/296) of S-healthy *sequences* (Figure S11b). Therefore, dbBact hints at a common gut response to acute pancreatitis, diarrhea, and Crohn’s disease, i.e., a phenomenon of general dysbiosis formerly suggested by Duvallet et al. (20). Next, we focused on the twelve *sequences* in S-panc that are not associated with either Crohn’s disease or ulcerative colitis across dbBact *annotations*. The *terms* associated with these thirteen *sequences* include “fermentation,” “homo sapiens,” “feces,” and “skin,” as well as “soil” and *sequences* tagged as “candidate contaminants” (Figures S11c-d). Additional experimental validation is required to determine whether some of these bacteria are pancreatitis-specific.

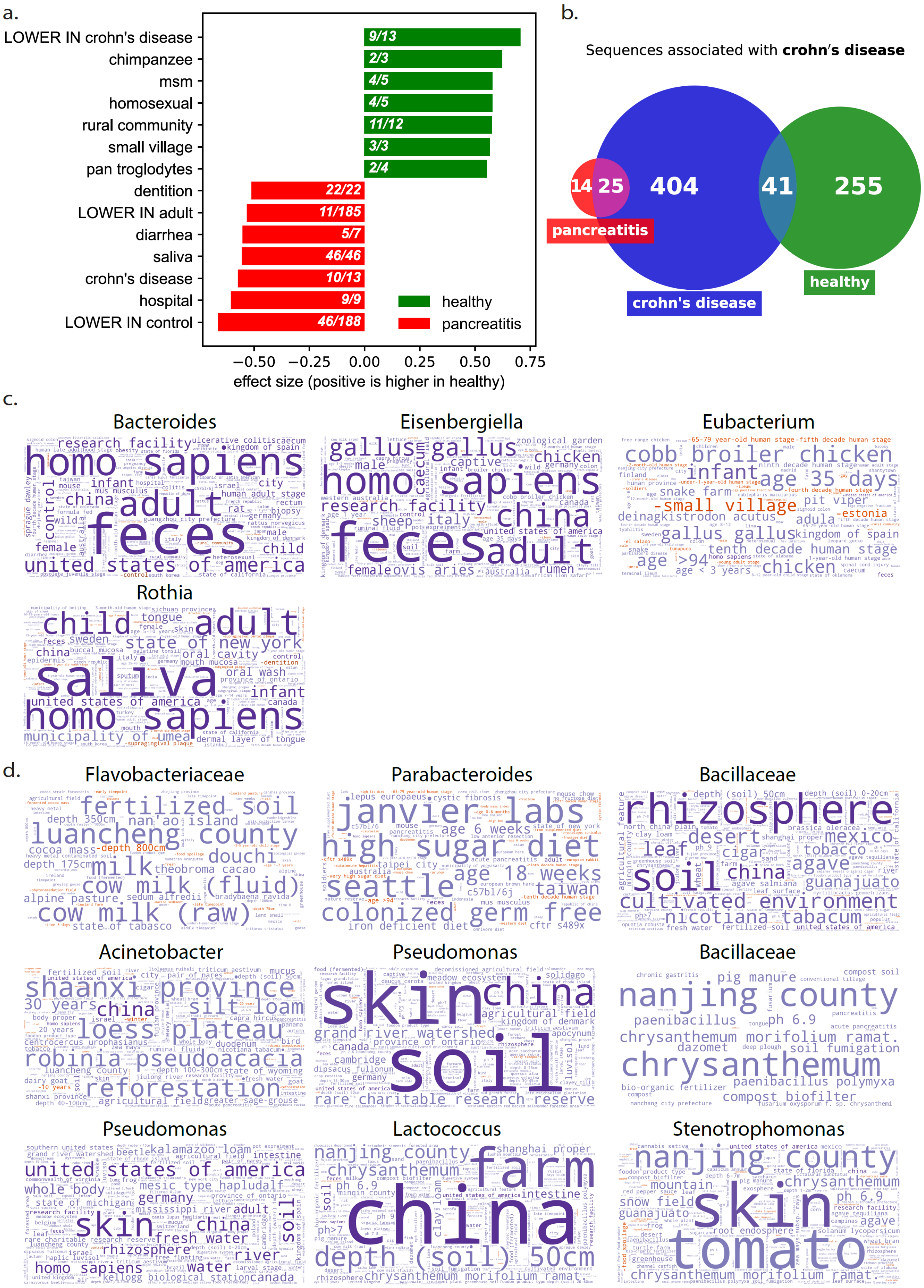
a.
LOWER IN crohn's disease
chimpanzee
msm
homosexual
rural community
small village
pan troglodytes
dentition
LOWER IN adult
diarrhea
saliva
crohn's disease
hospital
LOWER IN control
9/13
2/3
4/5
4/5
11/12
3/3
2/4
22/22
11/185
5/7
46/46
10/13
9/9
46/188
healthy
pancreatitis
−0.50 −0.25 0.00 0.25 0.50 0.75
effect size (positive is higher in healthy)
b.
Sequences associated with **crohn's disease**
14
25
404
41
255
pancreatitis
crohn's disease
healthy
c.
Bacteroides
Eisenbergiella
Eubacterium
Rothia
d.
Flavobacteriaceae
Parabacteroides
Bacillaceae
Acinetobacter
Pseudomonas
Bacillaceae
Pseudomonas
Lactococcus
Stenotrophomonas

**Figure S11: Acute pancreatitis and Crohn's disease share gut bacteria. a.** Bar plot of enriched *terms* in fecal samples of acute pancreatitis vs. healthy controls. **b.** Venn diagram for the *term* "crohn's disease." **c-d.** The list of 40 *sequences* higher in pancreatitis patients than in healthy controls was pruned to include *sequences* that are not associated with either Crohn's disease or ulcerative colitis across dbBact *annotations*, resulting in 13 bacteria whose word clouds are shown (see Supplementary File 6 for the sequences). **c.** The *terms* associated with 10 of these *sequences* include bacteria related to fermentation (Flavobacteriaceae), human feces and skin (Bacteroides), and soil (Bacillaceae). **d.** Three of the 13 *sequences* were annotated in dbBact as candidate reagent-related contaminants.

##### Chronic fatigue syndrome patients are enriched in bacteria observed in people with little physical activity

Giloteaux et al. (21) collected samples from 48 chronic fatigue syndrome (CFS) patients and compared them with those of 39 healthy controls. dbBact standard analysis identified 19 *sequences* higher in CSF patients, and 40 *sequences* higher in controls, referred to as S-CSF and S-Healthy, respectively. *Sequences* that were more abundant in the CFS group are enriched in *terms* related to low physical activity, while healthy controls are enriched in *terms* related to rural and less industrialized communities (Figure S12a).

For example, 27% (11/40) of control *sequences* are associated with the *term* "physical activity," compared to 0% (0/19) of CSF-related *sequences* (Figure S12b). Analogously, 68% (13/19) of S-CFS *sequences* are shared with "little physical activity" *sequences* across dbBact, compared to 0% of S-Healthy *sequences* (Figure S12c). Note that these associations are derived from a single dbBact experiment, which is based on the American Gut project data (15). Two possible interpretations for this observation are: (i) the bacterial difference in CFS patients is due to lower physical activity, and hence observed also in American Gut participants with lower physical activity; (ii) the microbiome change in CFS is due to the disease rather than to the physical activity level, and the associations with American Gut are due to unreported CFS patients in the American Gut cohort.

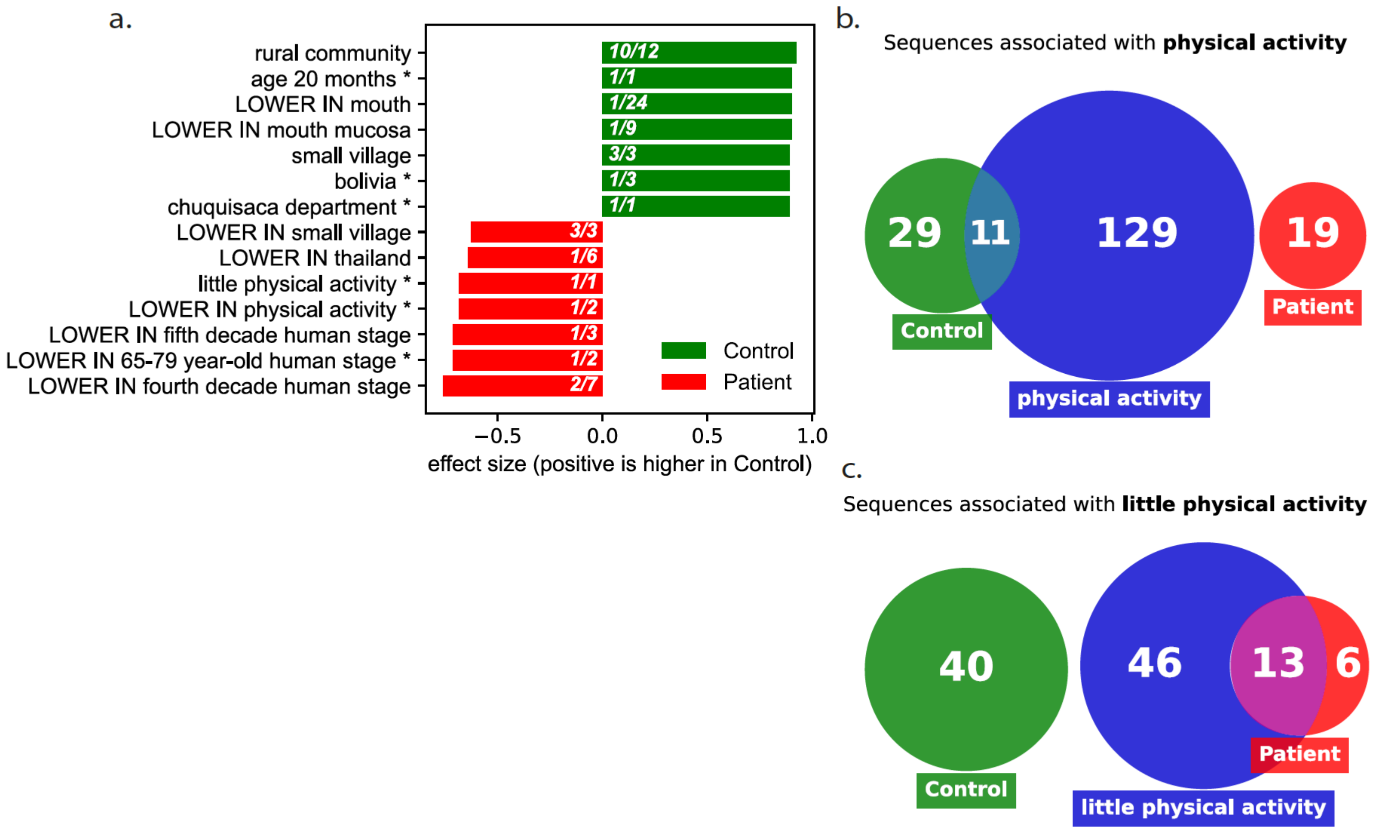


**Figure S12: Chronic Fatigue Syndrome (CFS) patients are enriched in bacteria observed in people with little physical activity**. a. Bar plot of enriched dbBact *terms* comparing the *sequences* higher in healthy controls (green) or CFS patients (red). b. Venn diagram showing the number of *sequences* associated with dbBact *annotations* containing the *term* "physical activity." Green and red circles represent the *sequences* higher in healthy controls and CFS patients, respectively, and the blue circle corresponds to all dbBact *sequences* associated with the *term* "physical activity." c. Same as b for the dbBact *term* "little physical activity."

##### Bank voles inhabiting regions inside and outside the Chernobyl Exclusion Zone: Differences in skin microbial communities may be attributed to exposure to humans and farm animals rather than to radioactivity

Lavrinienko et al. collected skin swabs of bank voles, *Myodes glareolus*, inside the uninhabited Chernobyl Exclusion Zone (CEZ), and in the outskirts of Kyiv, Ukraine, i.e., outside the contaminated region (22). 110 samples were collected from five sites within the CEZ, having different levels of environmental radioactivity, and 46 samples were collected in two locations around Kyiv. Standard analysis resulted in 1,203 and 327 *sequences,* more abundant inside and outside the CEZ, respectively. These sequences, referred to as S-CEZ and S-Kyiv, were submitted to dbBact as an enrichment query. The bar plot in Figure S13a displays overrepresentation of human and farm animal *terms* in S-Kyiv, as opposed to plant-related bacteria in S-CEZ. To validate this finding, we compared S-CEZ *sequences* with all 9,233 dbBact *sequences* having an *annotation* of "plant;" 38% (453/1203) of S-CEZ bacteria were independently assigned a "plant" *annotation* across dbBact, as opposed to only 18% (58/327) of S-Kyiv *sequences* (Figure S13b). It is reasonable to hypothesize, therefore, that at least some of the differences in skin microbial communities of bank voles inhabiting the two regions are not due to radioactivity but may be attributed to differences in exposure to humans and farm animals or their byproducts (which are less common in the CEZ).

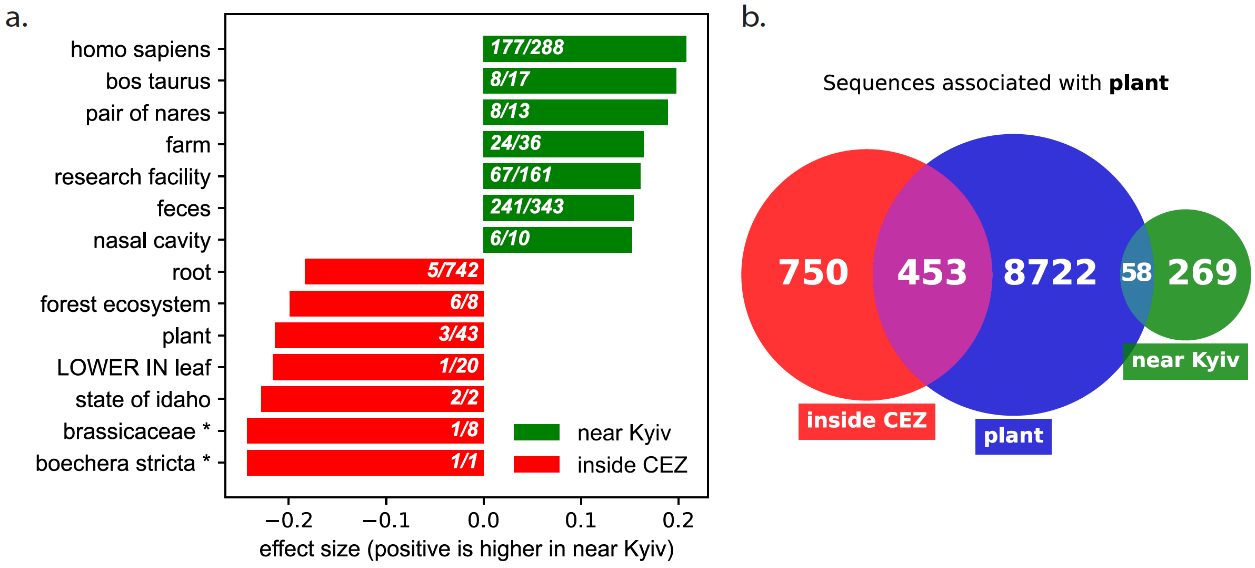


**Figure S13: Bank voles inhabiting regions inside and outside the Chernobyl Exclusion Zone: Differences in skin microbial communities may be attributed to exposure to humans and farm animals rather than to radioactivity**. **a.** Bar plot of enriched *terms*. **b.** Venn diagram for "plant" displaying the number of dbBact *sequences* associated with the *term* in either group.

#### Diurnal oscillations in meerkat feces are driven by soil bacteria

Risely et al. (23) collected over a thousand fecal samples from South African wild meerkats (*Suricata suricatta*), observing strong diurnal oscillations in microbiome composition. They found that morning and afternoon samples were significantly different, eclipsing seasonal and lifetime dynamics.

Using standard analysis, we looked for differentially abundant bacteria between the morning and afternoon samples, identifying 568 *sequences* higher in morning samples, and 4400 *sequences* higher in afternoon samples. dbBact *term* enrichment indicates enrichment of soil-related *terms* in afternoon samples (e.g., "soil," "rhizosphere," "desert," "triticum aestivum") (Figure S14a). Examining the number of *sequences* in each group that are associated with the *term* "soil" shows that 29% (1,268/4,384) of afternoon-related *sequences* are associated with "soil," compared to only 3% (20/567) of morning-related *sequences* (Figure S14b). To further illustrate this effect, Figure S14c displays the fraction *sequences* associated with the *terms* "soil" and "feces" in the course of the day.

The fraction of "soil" *sequences* rises from 15% in the morning to about 30% in the afternoon, while the fraction of "feces" displays an opposite trend. Our results show that soil-associated bacteria are driving the diurnal oscillations in meerkat feces.

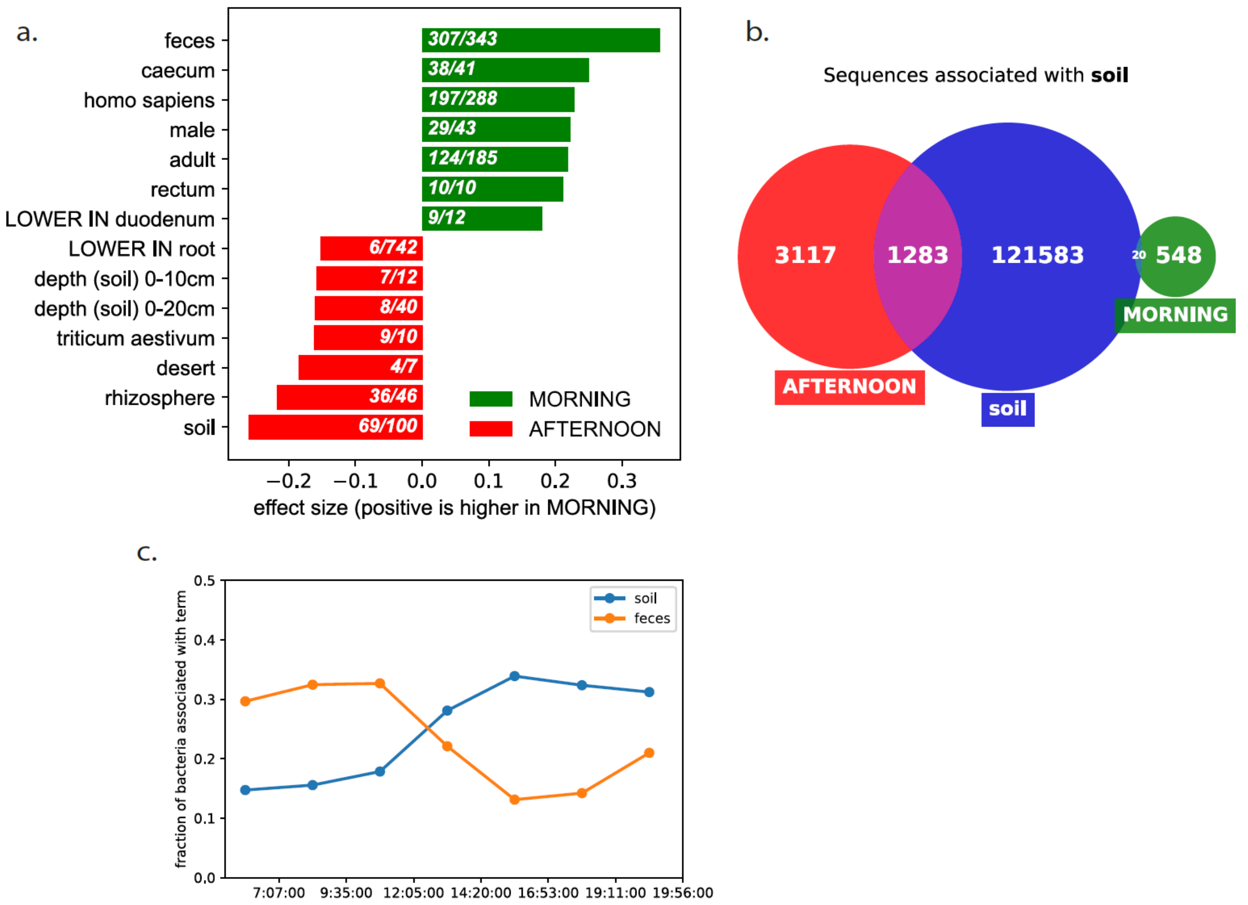


**Figure S14: Soil/feces origin of diurnal oscillation in wild meerkats. a.** Bar plot of enriched dbBact *terms* comparing the *sequences* higher in morning samples (green) and afternoon samples (red). **b.** Venn diagram showing the number of *sequences* associated with dbBact *annotations* containing the *term* "soil." Green and red circles are the *sequences* higher in morning and afternoon, respectively, and the blue circle corresponds to all dbBact *sequences* associated with the *term* "soil." **c.** Fraction of *sequences* associated with "soil" or "feces" (blue and orange, respectively) in samples collected along the day. We split the samples into bins of 2.5 hours and collected the list of sequences that appeared in at least 10% of the bin's samples. *Sequences* were then classified as feces-associated in case their *annotations* contained the term "feces." An analogous classification was performed for soil-associated *sequences*. If the *annotations* of a *sequence* contained both "feces" and "soil," its classification was based on the highest $F_1$ score.

#### Tracking sources of airborne bacteria: Clear days display fecal bacteria from human and farm animals, whereas dust storms carry desert and soil associated bacteria

Gat et al. profiled air samples in an urban region (Rehovot, Israel) during a dust storm and during clear days (24). Our re-analysis detected 418 and 182 *sequences* as significantly enriched during dust storm and clear conditions, respectively. These sequences, referred to as S-dust and S-clear, were submitted to dbBact as an enrichment query (Figure S15a). Enriched *terms* in S-dust include "'soil" and "desert," as opposed to "homo sapiens," "feces," and other anthropogenic-like *terms* in S-clear. To validate this finding, we compared S-dust *sequences* with all 6,310 dbBact *sequences* having an *annotation* of "desert;" 66% (271/418) of S-dust bacteria were independently assigned a "desert" *annotation* across dbBact, as opposed to only 18% (33/182) of S-clear *sequences* (Figure S15b). An analogous analysis performed over "homo sapiens" and "feces" identified 61% and 55%, respectively, of S-ambient sequences in other dbBact experiments, compared to only 18% and 8%, of S-dust sequences (Figure S15c,d). Thus, in this case, dbBact revealed the sources of the samples: fecal bacteria from human and farm animals are airborne during ambient weather conditions, whereas dust storms bring over desert and soil associated bacteria.

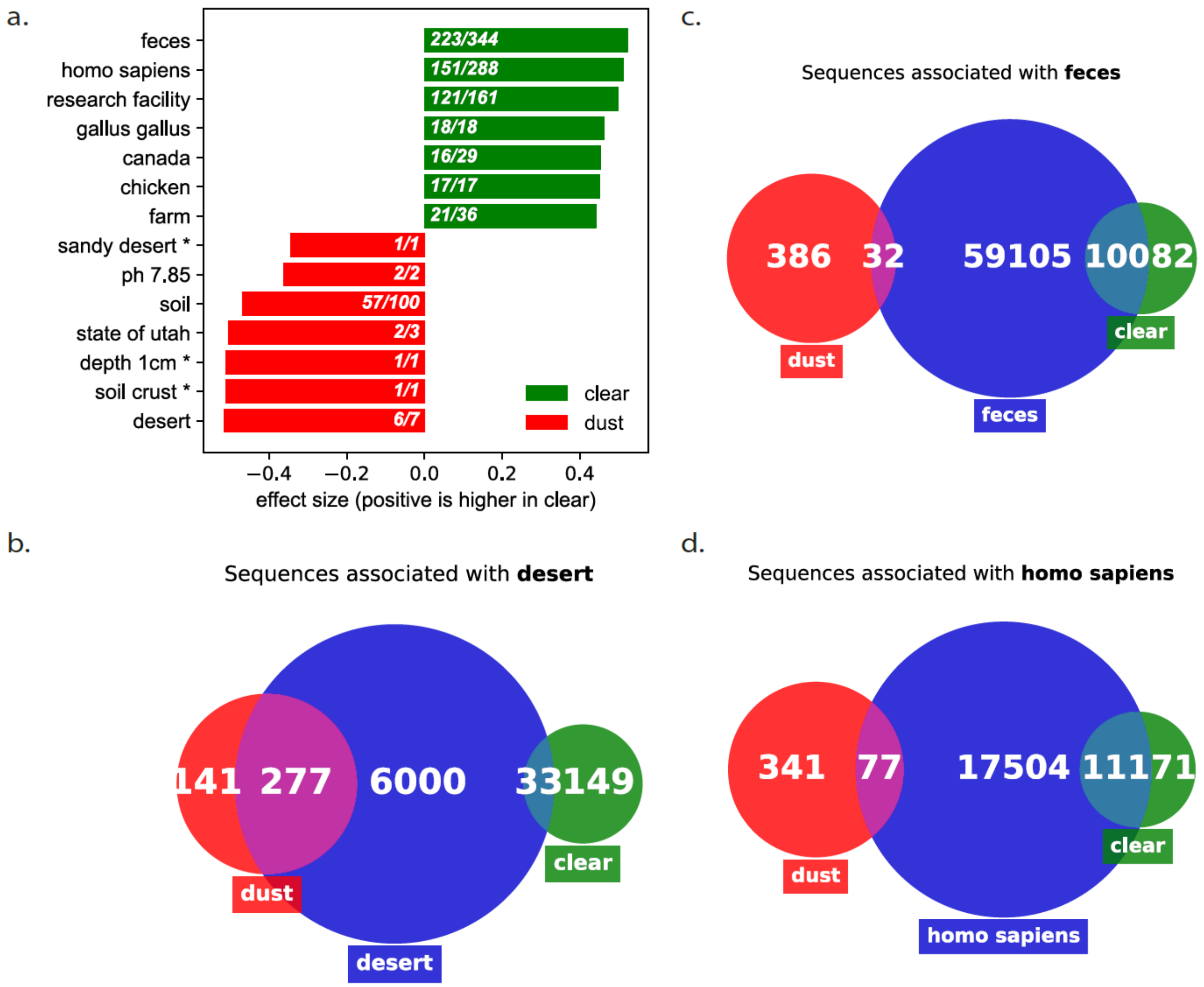


**Figure S15: Tracking sources of airborne bacteria: Clear days display fecal bacteria from human and farm animals, whereas dust storms carry desert and soil associated bacteria. a.** Bar plot of enriched *terms*. **b.** Venn diagram for "desert" displaying the number of dbBact *sequences* associated with the *term* (blue circle), and their overlap with *sequences* associated with "desert" in either group (red and green circles). The numbers indicate the number of *sequences* in each part of the Venn diagrams. **c.** and **d.** Similar Venn diagrams for "feces" and "homo sapiens," respectively.

#### Microbiome composition along the Bronx River is affected by salinity due to proximity to the ocean

Naro-Maciel et al. (25) collected water samples in two locations, Hunts Point and Soundview Park, along the Bronx river in New York (Figure S16a). Using standard analysis, we detected *sequences* differentially enriched in either location, resulting in 30 and 106 *sequences* higher in Hunts Point and Soundview Park, respectively. *Terms* enriched in Hunts Point are related to freshwater (e.g., “fresh water,” “lake,” “river”), while Soundview Park is enriched in ocean water *terms* (e.g., “ocean,” “pacific ocean,” “sea water”) (Figure S16b). These results indicate that although both locations harbor a mainly saline water-related microbiome (data not shown), Hunts Point, which is farther from the ocean, is enriched in fresh water-related bacteria.

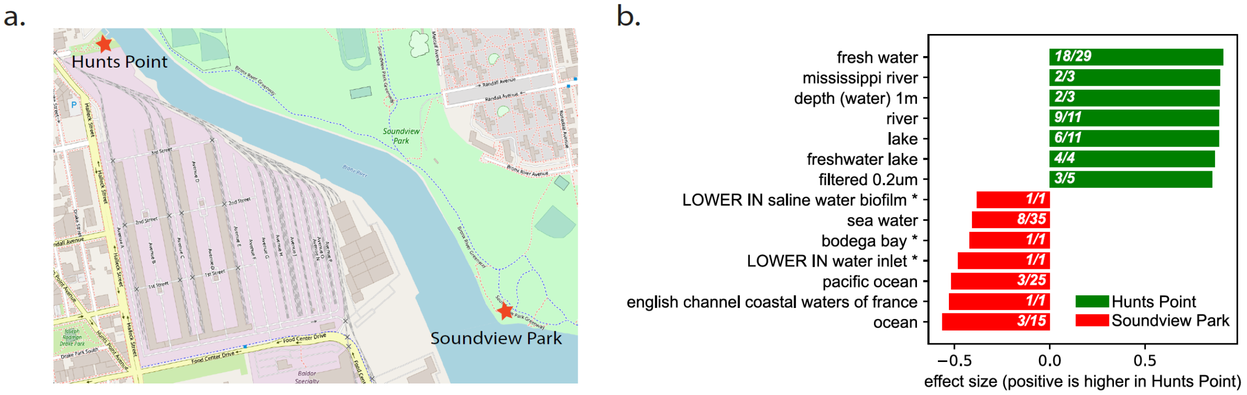


**Figure S16: Microbiome composition along the Bronx River is affected by ocean proximity. a.** Map showing the sampling locations (star) of Hunts Point and Soundview Park along the Bronx River, New York. **b.** Bar plot of the top 10 enriched dbBact *terms* comparing the *sequences* higher in Hunts Point (green) and Soundview Park (red).

#### Fecal samples from toilet paper can contain significant amounts of skin bacteria

Caporaso et al. followed the oral, skin, and fecal microbiome of an individual using daily samples for one year. dbBact *term*-based PCA (see Methods) shows a separation between oral, fecal, and skin samples (Figure S17a) along two principal axes whose interpretation is provided below.

*1st principal axis (horizontal) - feces vs. saliva*: the *terms* with the highest positive coefficients in the 1st principal component are "feces," "LOWER IN crohn's disease" and "LOWER IN ulcerative colitis," while the *terms* with the most negative coefficients are "saliva," "mouth," "LOWER IN supragingival plaque." Hence, the horizontal axis is the "feces vs. saliva" principal axis, where higher values correspond to feces. Indeed, most fecal samples (blue) have high values along this axis whereas most saliva samples (red) have low values.

*2nd principal axis (vertical) - skin vs. feces/saliva*: The *terms* with the highest positive coefficients in the 2nd principal component are "skin," "pair of nares" and "nasal cavity," while the most negative coefficients are "homo sapiens," "adult," "feces" and "saliva." Hence the vertical axis is the "skin vs. feces/saliva" axis.

While saliva samples form a tight cluster at the bottom left, fecal samples show a spread along the 2nd principal axis. To further investigate this behavior, we display the dbBact *term*-based PCA of only fecal samples (Figure S17b). The 1st principal axis in this case (which holds 60% of the variance) is a "feces vs. skin/vagina" axis. Some samples are spread along this axis, with a majority of samples located towards the "feces" direction of the axis (blue circles), whereas a smaller group is located towards the "skin"/"vagina" direction (magenta circles), indicating a possible skin-derived contamination in this set of samples. To validate this, Figure S17c displays a heatmap of the fecal samples sorted according to sampling date, where the top color bar corresponds to blue or magenta samples in Figure S17b, and each *sequence* is "classified" according to its largest $F_1$ score *term* out of "feces" and "skin." As can be seen, the samples originating from the magenta samples contain a cluster of skin-associated *sequences*. This agrees with sporadic contaminations of fecal samples with skin-associated bacteria in some of the samples. As the sampling protocol involved swabbing of used bathroom tissue, this raises the possibility that skin bacteria from the rectum were sometimes sampled from the tissue paper, leading to the appearance of this skin-related cluster.

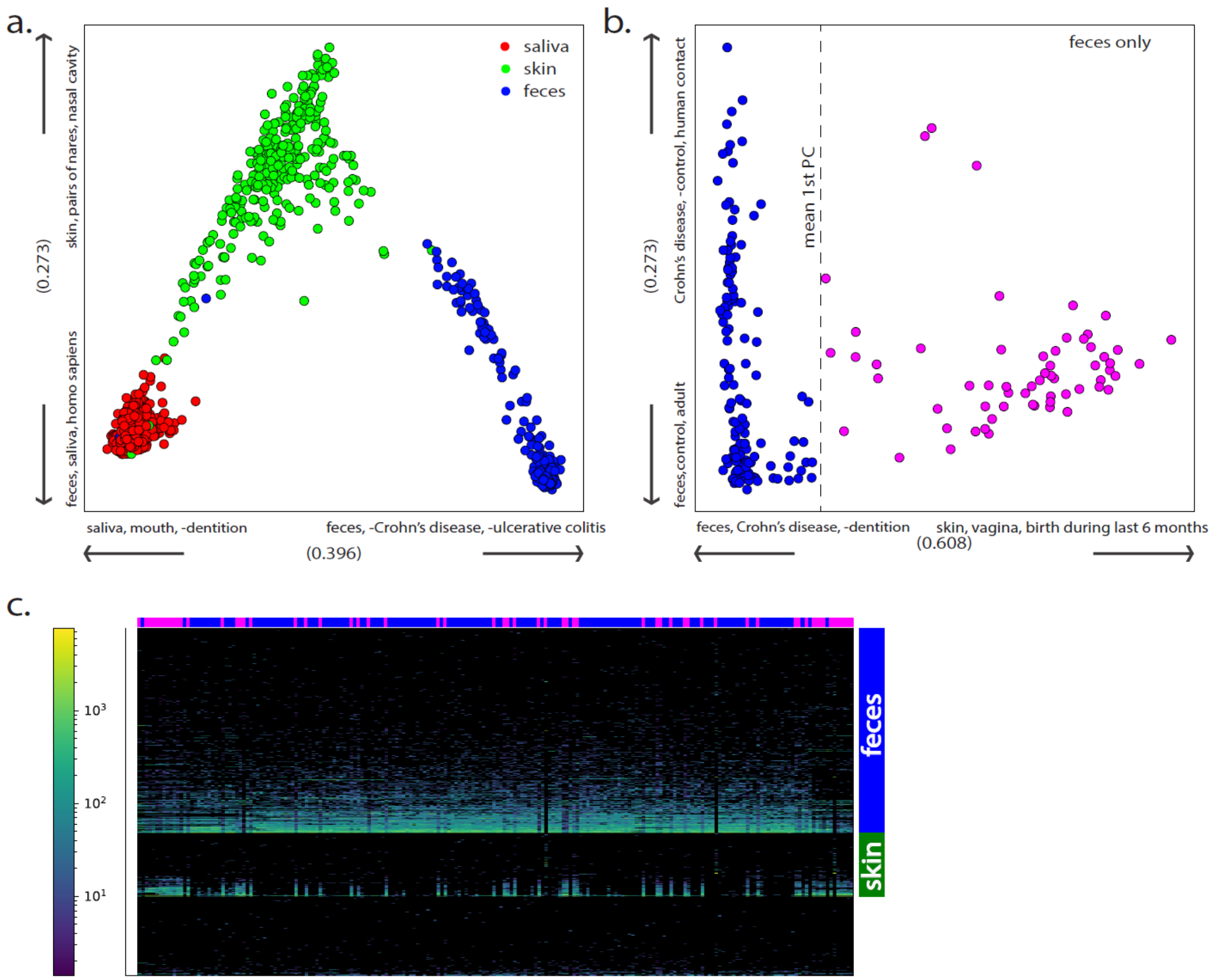


**Figure S17: Fecal samples from toilet paper can contain significant amounts of skin bacteria. a.** dbBact *term*-based PCA of all samples from one individual. Frequency weighted *term* precision scores are used to construct the distance matrix (see Methods section for details). The three top terms contributing to each axis direction are shown. Variance explained is 0.4 and 0.27 for the first and second axes, respectively. **b.** dbBact *term*-based PCA for only fecal samples colored by their 1st PCA coordinate (blue and magenta for projections lower or higher than the mean value, respectively). **c.** Heatmap showing the frequency of each *sequence* (row) in consecutive daily fecal samples of the same individual (columns). Each sequence is classified according to the *term* with the highest $F_1$ score out of the terms "feces" and "skin." The horizontal color bar denotes samples belonging to the blue and magenta groups in (b).

#### Filtering mouse-associated contaminants in human nasopharyngeal samples improves statistical power

Xu et al. analyzed the nasopharyngeal microbiome of six- and twelve-months old infants, aiming to study the effect of *Streptococcus pneumoniae* colonization (27). Examining the word cloud of the *terms*, we observed several seemingly unrelated *terms*, e.g., "mouse," "mus musculus" and "soil" (Figure S18a). Therefore, we "classified" each bacterium in this study according to its highest $F_1$ score *term* out of "homo sapiens," "mus musculus," "soil," and "other." The resulting heatmap shows a cluster of mouse-associated *sequences* that appear together in a subset of the samples, as well as a small set of soil-associated *sequences* (Figure S18b). These findings indicate a possible contamination, maybe due to sample processing or reagent-borne contaminants. Because these bacteria are spread across sample types, they do not introduce a systemic bias in the authors' findings. But such contamination may reduce the power of downstream statistical tests. Therefore, we removed all mouse- and soil-associated *sequences* and reapplied total sum per sample (TSS) normalization. Consequently, the difference in Shannon diversity between the six- and twelve-months age groups, which was not statistically significant before contamination filtering (Mann-Whitney non-parametric test p-value=0.16, Figure S18c), became significant (Mann-Whitney non-parametric test p-value=0.049, Figure S18d).

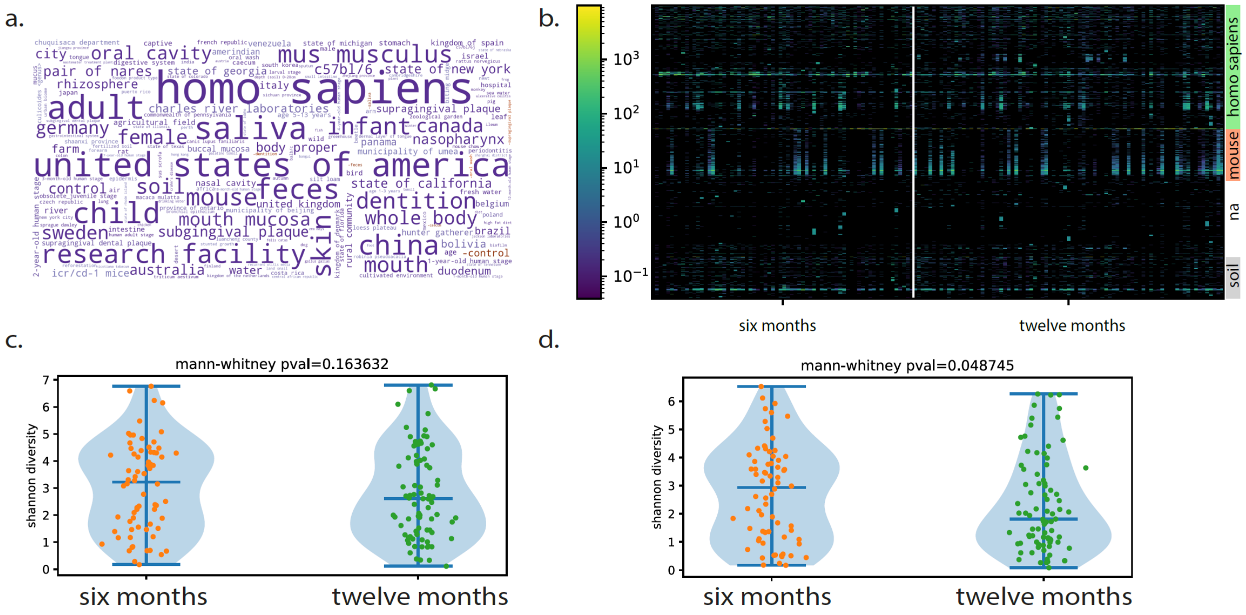


**Figure S18: Detection and removal of contaminants increases statistical power**. **a.** dbBact *term* word cloud of prevalent *sequences* (present in >0.3 of samples) of infant nasopharyngeal samples taken at age six and twelve months. **b.** Heatmap showing the frequency of each *sequence* (row) in the infant nasopharyngeal samples (columns). Each *sequence* is classified according to the *term* with the highest $F_1$ score. **c-d.** Comparison of the Shannon diversity between the six- and twelve-months age groups without (left) and with (right) filtering *sequences* associated with "mus musculus" or "soil."

### Supplementary methods

#### dbBact - Implementation

dbBact data are stored in a SQL database using Postgres 9.5.10. A weekly dump of the complete dbBact database (excluding users' private details) is available at https://dbbact.org/download

#### Database tables

The basic structure of the database is shown in Figure S19. A full database schema is shown in Figure S20, and a detailed list of all dbBact tables and columns is available in Supplementary File1 (database-tables.xls). General implementation notes are provided below.

*Sequences* (dbbact.SequencesTable): A *sequence* is a partial 16S rRNA sequence of at least 100nt length. Because different *experiments* can have different read lengths, the same bacterium may be described by multiple *sequence* entries of different lengths. Similarly, because *experiments* may amplify different regions, the same bacterium may be represented by multiple *sequences* originating from different regions. dbBact assigns a taxonomy to each uploaded *sequence* based on RDP (28) and Greengenes 13.8 (29), using an external script applied daily.

*Experiments* (dbbact.ExperimentsTable): The table holds a list of the different *experiments* available in dbBact. The *experiments* describe the source of the dataset (i.e., the SRA/qiita (30) accession, the DOI, the title of the paper, etc.).

*Terms* (dbbact.OntologyTable): dbBact *terms* are ontology based, resulting in several tree structures stored in dbbact.OntologyTreeStructureTable. Users may also add new *terms* to dbBact, if needed.

*Annotations* (dbbact.AnnotationsTable): The different *annotation* predicates (Table S3) are stored in dbbact.AnnotationTypesTable. Each *annotation* is based on a single dataset, but several *annotations* can originate from the same dataset (e.g., an *experiment* containing sick and healthy subjects can be assigned *annotations* describing the common *sequences* in either the sick and healthy groups, as well as *annotations* describing *sequences* higher in the sick group than in the healthy group, and *vice versa*). Each *annotation* is associated with the *experiment* it is derived from and with the user who added the *annotation*.

Associations between an *annotation* and its *sequences* are stored in dbBact.SequencesAnnotationTable. *Terms* in each *annotation* appear in dbbact.AnnotationListTable.

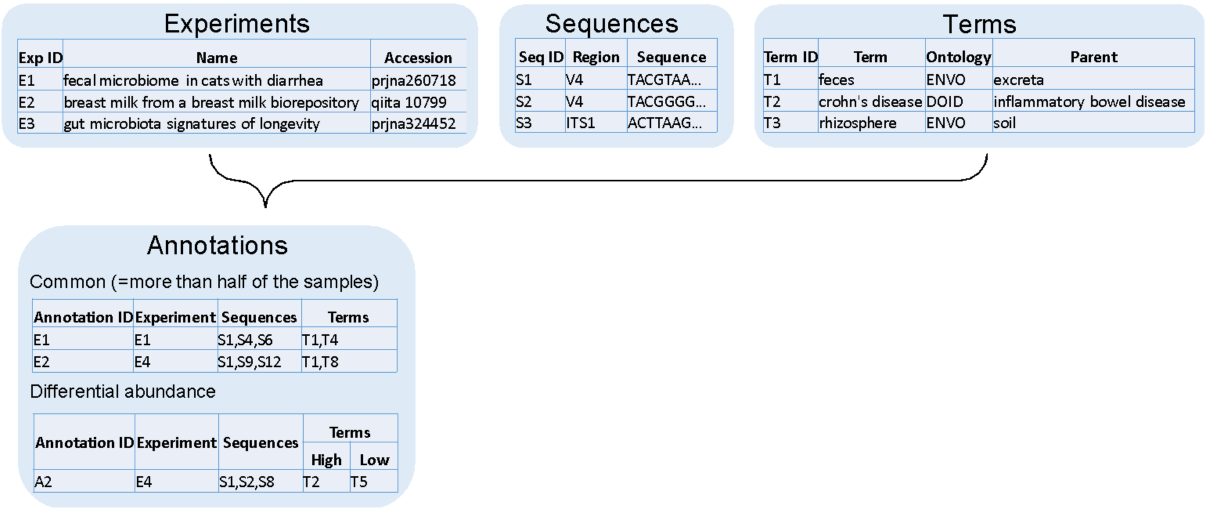


**Figure S19: Main dbBact entities.** Each *annotation* associates several bacterial *sequences* with a set of ontology *terms* describing various phenotypes based on an *experiment*.

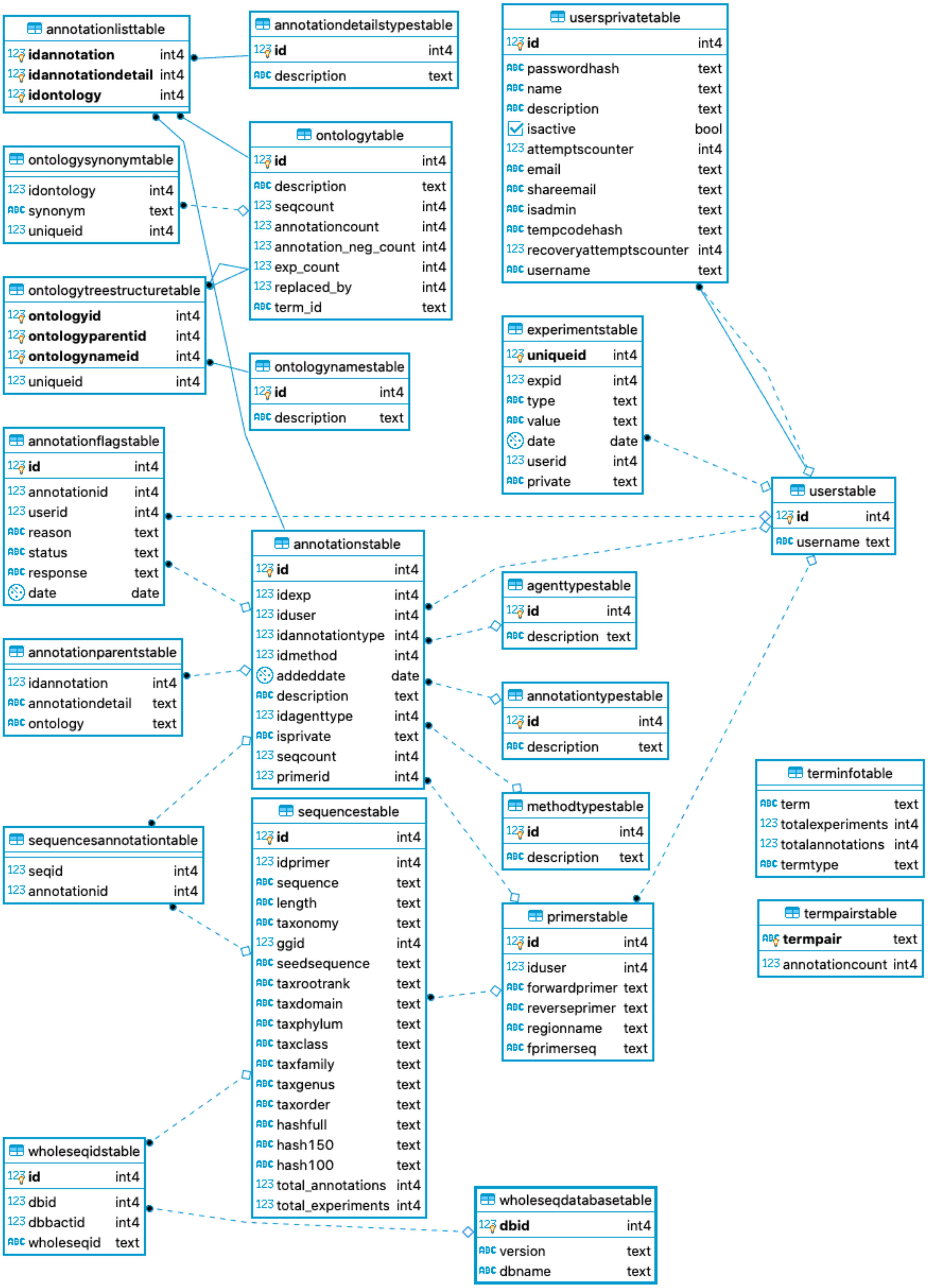


Figure S20: dbBact database schema.

#### Ontologies

Table S1 presents ontologies available in dbBact release 2021.05. In addition, new *terms*, which do not appear in either ontology, can be added to the generic dbBact ontology. dbBact stores each ontology *term* as a directed graph, with parent *terms* defined as *terms* appearing in the ontology, like one of the following:

'is_a'/'derives_from'/'located_in'/'part_of' /'develops_from'/'participates_in'.

| Ontology | Version |
|---|---|
| ENVO (Environment Ontology) (1,2) | 2019-03-14 |
| DOID (Human Disease Ontology) (31) | 2019-09-16 |
| EFO (Experimental Factor Ontology) (32) | 3.10.0 |
| GAZ (Gazetteer) | 2013-12-23 |
| HSAPDV (Human Stages Ontology) | 2018-05-20 |
| PATO (Phenotype and Trait Ontology) | 2019-12-03 |
| TO (Plant Trait Ontology) (33) | 2019-05-21 |
| UBERON (Uber-Anatomy Ontology) (5) | 2018-11-25 |
| NCBI Taxonomy (4) | 2019 |

**Table S1**: List of ontologies supported in dbBact release 2021.05.

#### Primer pairs currently implemented in dbBact

| Region | Forward primer (dbBact sequences start at the end of the primer sequence) |
|---|---|
| V1-V2 | AGAGTTTGATC[AC]TGG[CT]TCAG |
| V3-V4 | CCTACGGG[ACGT][CGT]GC[AT][CG]CAG |
| V4 | GTGCCAGC[AC]GCCGCGGTAA |

**Table S2**: **Primers used for dbBact *sequences***. Square brackets denote nucleotide degeneration (e.g., [AG] denotes A or G at this position).

#### Predicates used in *annotations*

| Predicate | Description | Example |
|---|---|---|
| COMMON | Present in over half the samples of a given type in the *experiment* | COMMON in feces, homo sapiens, adult, State of Colorado |
| DOMINANT | Mean frequency >1% in samples of a given type in the *experiment* | DOMINANT in desert, rhizosphere, agave desert, State of California |
| HIGH/LOW (differential abundance) | Significantly different between two conditions in the *experiment* | HIGH in Crohn's disease compared to control in feces, homo sapiens, child, United States of America |
| CONTAMINANT | Suspected as a contaminant in an *experiment* | CONTAMINANT |
| OTHER | Additional observations (i.e., known pathogen, free text descriptions, etc.) | OTHER: Tropheryma whipplei (pathogen, whipple disease, whipple's disease)<br><br>OTHER: associated with glucose tolerance (mus musculus, caecum, feces, c57bl/6j, glucose tolerance) |

**Table S3: Available predicates in dbBact**, i.e., "relations" between dbBact *annotation* types.
